# Infusible Extracellular Matrix Biomaterial Enhances Cell-Specific Pro-Repair Responses Following Acute Myocardial Infarction

**DOI:** 10.1101/2025.11.08.687255

**Authors:** Joshua M. Mesfin, Alexander Chen, Quincy P. Lyons, Michael B. Nguyen, Maria L. Karkanitsa, Justin Yu, Van K. Ninh, Emily Gardner, Elyse G. Wong, Sriya N. Paleti, Julian Cheng, Benjamin D. Bridgelal, Kate E. Reimold, Selena Cao, Connor Uhre, Colin G. Luo, Zhenxing Fu, Kevin R. King, Karen L. Christman

## Abstract

To mitigate the pathological effects of myocardial infarction, we developed and investigated pro-reparative decellularized extracellular matrix (ECM) biomaterials: an intravascularly infusible ECM (iECM). However, the cellular and molecular mechanisms by which iECM mediates repair are unknown because investigations have relied on bulk techniques. Here, we leverage single nucleus RNA sequencing (snRNAseq) to measure pro-repair in various cell types across acute timepoints (1, 3 and 7 days post infusion). In iECM, we found pro-reparative macrophage activation, fibroblast remodeling, increased vasculaure development, lymphangiogenesis, cardioprotection, and neurogenesis. These findings were validated through spatial transcriptomics. Thus, we define the pro-reparative nature of decellularized ECM biomaterials on cardiac cell types and elucidate previously undiscovered therapeutic pathways, further demonstrating the potential of iECM as an MI therapy as well as display the wealth of data generated from next-generation sequencing.

## 1. Introduction

Acute myocardial infarction (AMI) is characterized by occlusion of coronary arteries which deprives the dependent myocardium of oxygen-rich blood^[1]^. This leads to a molecular and cellular cascade of ischemic injury, inflammation, cardiomyocyte apoptosis and necrosis, endothelial cell dysfunction and vascular leak, and fibroblast activation, which collectively contribute to negative left ventricular (LV) remodeling and clinical heart failure, a chronic progressive and ultimately fatal disease. The current standard of care for MI is percutaneous coronary intervention ^[1b, 2]^; however, there remains an unmet clinical need for therapies to limig negative LV remodeling post-MI and prevent heart failure, particularly those that are delivered minimally invasively.

AMI precipitates a vigorous inflammatory response that has been pursued as a target for therapy to improve outcomes^[3]^. It is characterized by a stereotyped response that begins with neutrophil infiltration, is followed by monocytes that differentiate into inflammatory and reparative macrophages, and finally lymphocytes. The ensuing inflammatory cascade post-AMI evolves in a time dependent manner. First, circulating neutrophils, monocytes, and lymphocytes infiltrate within 24 hours^[4]^. Monocyte infiltration is slower, peaking within 72 hours^[1b, 4]^. During this acute phase (<7 days), two distinct populations of macrophages contribute to either damage or healing^[4]^. This cascade leads to myocardial fibrosis and subsequent heart failure over the course of weeks and months^[1]^. Therefore, therapeutic intervention and delivery is desirable immediately upon reperfusion to target the inflammatory response.

In addition to the pathological immune response, MI also induces molecular cascades within non-immune cell types, such as cardiomyocytes, vascular endothelium, and fibroblasts. Immediately upon MI, at the site of ischemic injury, cardiomyocytes are known to undergo necrosis within the first day, with apoptosis continuing in the border zone post-MI^[5]^. Neuronal cells are more resilient post-ischemic injury, but also have reduced function. Upon injury, the vascular endothelium becomes permeabilized and remains leaky for up to 7 days post-MI^[6]^. Other vascular cell types, such as smooth muscle cells and lymphatic endothelial cells also undergo significant cellular death and limited proliferation^[7]^. In contrast, fibroblasts activate and proliferate starting from 1 days post-MI, peaking around 7 days^[6, 8]^. Given the limitations of conventional therapies, we and others have utilized injectable biomaterial therapeutics^[3, 9]^. We recently developed an intravascularly infusible decellularized extracellular matrix (iECM) material derived from porcine myocardium that can be delivered via intracoronary infusion upon reperfusion following percutaneous revascularization. This material localizes to leaky vasculature induced by AMI, reducing vascular permeability and exerting immunomodulatory and pro-reparative effects, which improve cardiac function^[5]^.

Our previous studies found that the iECM reduced cardiomyocyte apoptosis and increased vascular density as assessed by histology, and led to expression of genes associated with tissue repair and dampening of the post-MI immune response as determined by a custom Nanostring nCounter panel. However, as these findings utilized bulk sequencing techniques, understanding pro-repair requires more precise tools. Single nucleus RNAseq and spatial transcriptomics have revealed new post-MI mechanisms that may serve as new therapeutic targets^[8a, 10]^. We recently leveraged the technology to study potential mechanisms of action for pleiotropic biomaterials-based therapies locally administered to treat subacute and chronic models of MI^[11]^. Thus, to better understand the change in tissue response following iECM administration and thus mitigate the acute MI response, we leverage snRNAseq to uncover the temporal variation in the immune response, vascular healing, cardiomyocyte salvage, and fibroblast activation within the infarct post-AMI through pseudobulk analyses of relevant cell types. To validate our findings, we utilize spatial transcriptomics to score our transcriptional findings within the infarct of iECM treated samples to quantify the overall pro-reparative response.

## 2. Results and Discussion

### 2.1. Fabrication of iECM

To evaluate the therapeutic activity of iECM, a comparative study was conducted against a saline infusion in an AMI model. iECM was manufactured as previously described **(Figure 1A)** ^[12]^. Quality control characterization of the iECM via SDS-PAGE (Figure 1B), residual dsDNA content (Figure S1A), and viscosity (Figure S1B) were all consistent with prior reports, indicating reproducible manufacturing and a robust fabrication process^[12]^. To assess the effects of the iECM post-MI, iECM was delivered upon reperfusion using a simulated intracoronary injection in rats. To investigate the impact on the acute immune response, the timepoints of 1-, 3-, and 7-days post-MI were selected, corresponding to well-characterized phases of immune cell infiltration and function^[1b, 4]^. At each time point, heart sections were subject to nuclei isolation for RNA sequencing and spatial transcriptomics (Figure 1C).

**Figure 1.**
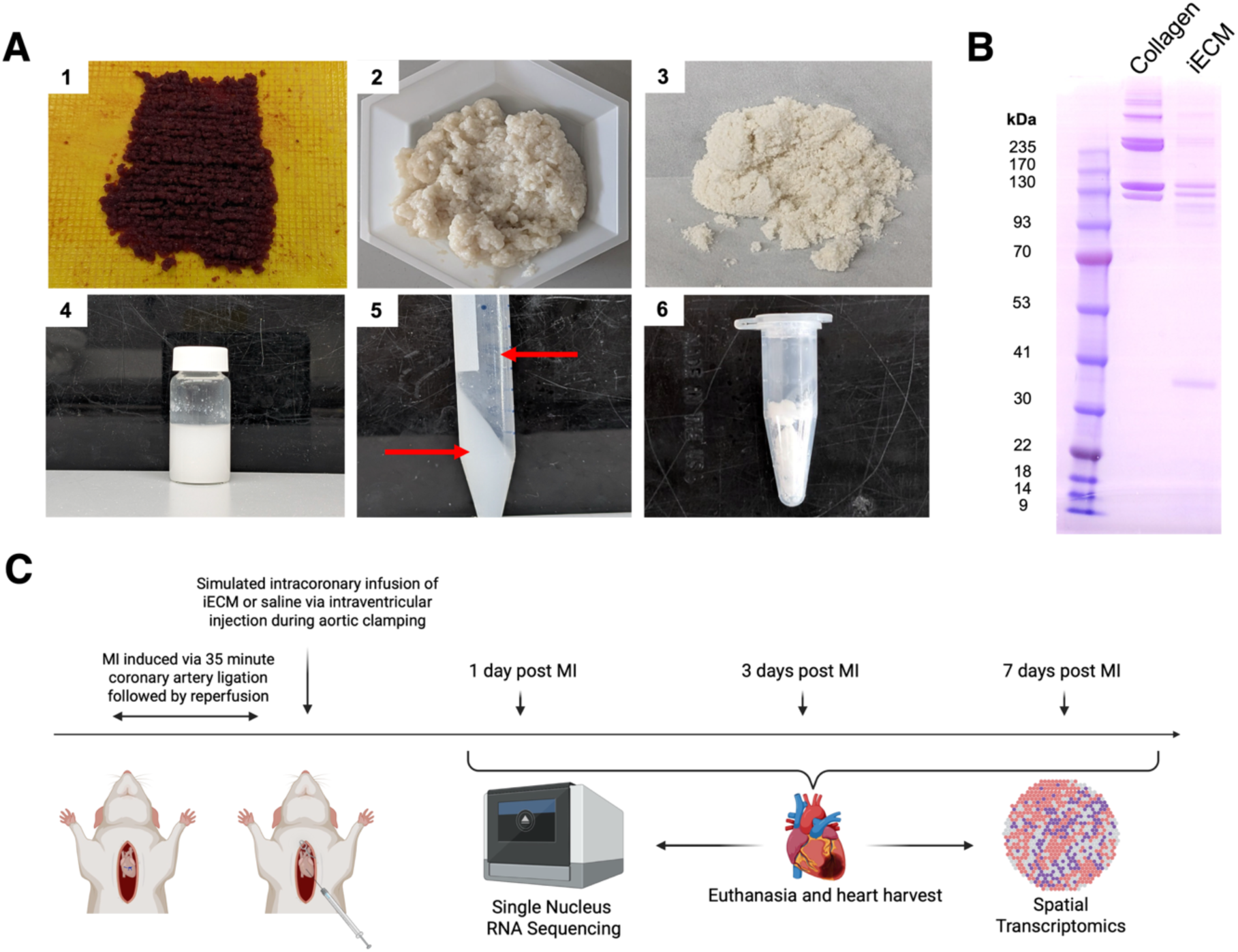
Infusible extracellular matrix (iECM) elicits downstream pro-repair activity post infusion in acute myocardial infarction (MI). A) Overview of iECM manufacturing, where porcine left ventricular myocardium is chopped into small pieces (1). The resulting ECM is washed with sodium dodecyl sulfate (SDS) (2) followed by rinsing, milled into a fine powder (3) and digested (4). High speed centrifugation is then performed to separate out large particulate matter (5) and is finally reconstituted for infusion for MI treatment (6). B) SDS-PAGE of iECM and collagen. C) Overall timeline for iECM bioactivity studies in acute MI. Simulated intracoronary infusion of iECM or saline was performed after MI and reperfusion. Hearts were harvested 1-, 3-, and 7-days post infusion. Samples were analyzed via single nucleus RNA sequencing (snRNAseq) and spatial transcriptomics.

### 2.2. Single Nucleus RNA Sequencing Captures Various Cell Types in Acute MI

For snRNAseq analysis, the features and counts were evaluated to remove cells below a quality threshold (Figure S2A, Figure S2B). The datasets across all timepoints were integrated and clustered, and we identified various cell types such as macrophages, endothelial cells, lymphatic endothelial cells, fibroblasts, mural cells, cardiomyocytes, and neuronal cells with the relative proportions of each cell type respective to treatment (Figures S2C-H). All identified cell populations are represented in the infarcted heart with marker genes already depicted in the literature^[8a, 13]^. Sample sizes (n = 4 biological replicates pooled into 2 sequencing replicates per condition except for day 3 iECM, which had n = 2 biological replicates and n = 1 sequencing replicate) and total number of cells captured are displayed in Table 1. The iECM treated samples were also integrated and clustered across time (Figure S2I-J) to investigate the changes in gene expression over time with respect to iECM treatment.

**Table 1.** Overview of Total Replicate and Nuclei/Spot Count for iECM Studies.

|  | **iECM**<br>**Day 1** | **Saline**<br>**Day 1** | **iECM**<br>**Day 3** | **Saline**<br>**Day 3** | **iECM**<br>**Day 7** | **Saline**<br>**Day 7** |
| --- | --- | --- | --- | --- | --- | --- |
| **snRNAseq**<br>**Replicates** | 4 biological replicates, pooled into 2 technical replicates | 4 biological replicates, pooled into 2 technical replicates | 2 biological replicates, pooled into 1 technical replicate | 4 biological replicates, pooled into 2 technical replicates | 4 biological replicates, pooled into 2 technical replicates | 4 biological replicates, pooled into 2 technical replicates |
| **Total**<br>**Nuclei** | 33661 | 44949 | 19526 | 64869 | 57048 | 56229 |
| **Spatial**<br>**Replicates** | 2 | 2 | 2 | 2 | 2 | 2 |
| **Total Spots** | 8345 | 6822 | 7496 | 7973 | 7906 | 7819 |

### 2.3. iECM Elicits Pro-Angiogenic Effects on Vascular Cells Earlier than Saline Treatment and is Preserved to Day 7

Previously, iECM has been shown to bind the leaky vasculature^[12a]^, a hallmark of areas of inflammation including ischemia reperfusion injury in MI. Thus, we first looked at endothelial cells to demonstrate how iECM localization to the leaky vasculature can lead to therapeutic effects on the underlying endothelium over time. As done previously^[11e]^, we subsetted and subclustered endothelial cells (*Flt1+/Ptprb+)* where we identified multiple endothelial cell populations **(Figure 2A)**, mapped the various phenotypes of the subpopulations found (Figure S3), and defined their relative proportions per treatment group (Figure S4A). All genes describing each subcluster are outlined in Table S1. To visualize the shift of the cells over time, the endothelial cell subclusters were integrated across treatments and timepoints. Uniform Manifold Approximation and Projections (UMAP) was used for dimensional reduction. The UMAPs were plotted, showing the shifts in the endothelial cell cluster (Figure 2A). The cells were then subsetted by timepoint for differentially expressed gene (DEG) analysis (Figure 2B) as well as analyzed via gene ontology (GO) pathway enrichment (Figure 2C). All differentially expressed genes for endothelial cells across time are outlined in Table S2.

**Figure 2.**
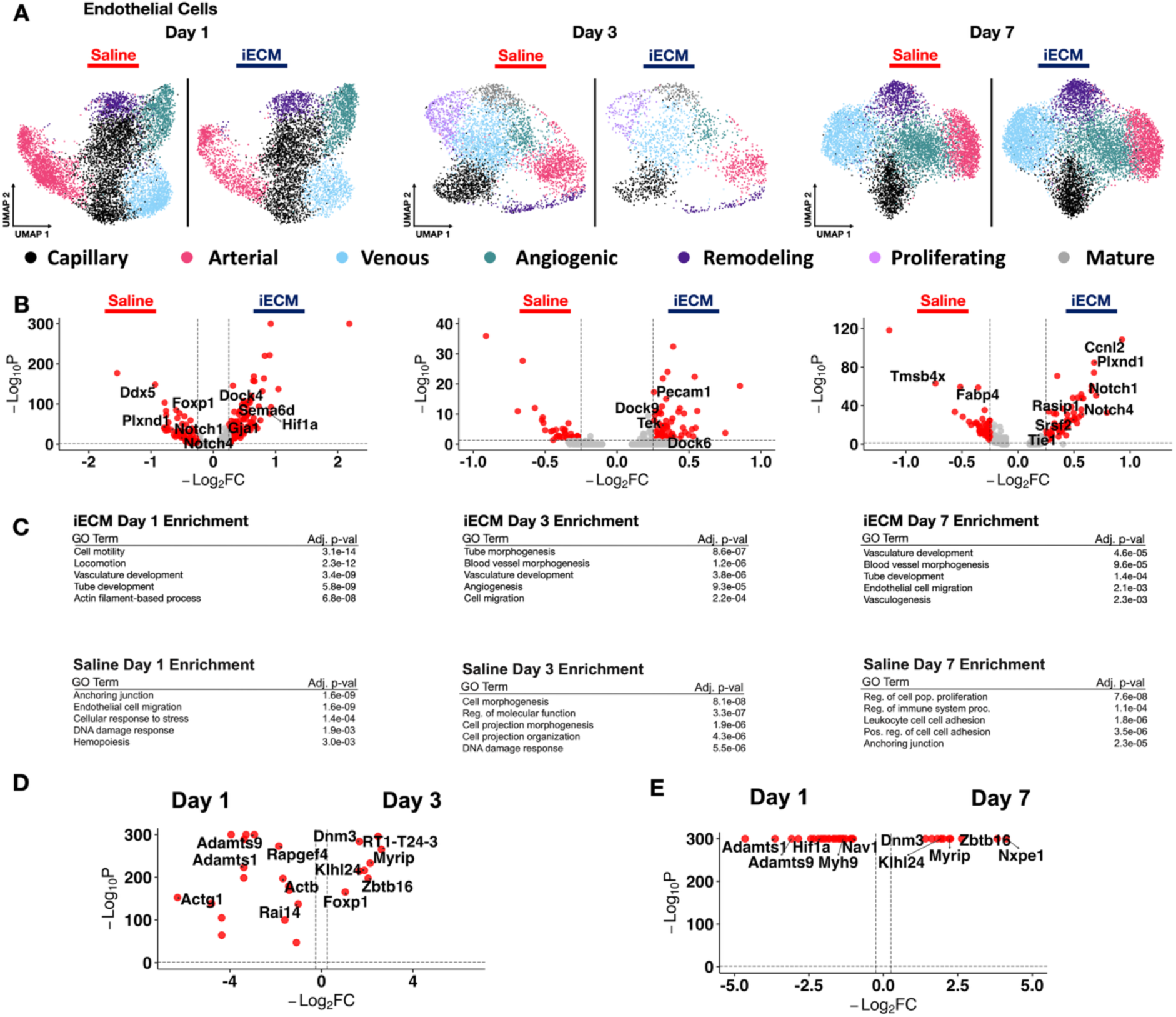
iECM promotes vasculature development at early timepoints. A) UMAP of endothelial cells in iECM and saline treated samples split across days 1, 3 and 7 post infusion. B) Differentially expressed genes in endothelial cells between iECM and saline at days 1, 3, and 7 post-infusion. C) All iECM specific differentially expressed genes were subjected to GO enrichment. D) Differentially expressed endothelial cell genes for iECM at day 1 versus day 3 post infusion. E) Differentially expressed endothelial cell genes for iECM at day 1 versus day 7 post infusion.

At day 1, we found that with iECM treatment, endothelial cells exhibit upregulation of genes such as *Hif1a*, which is implicated in mitochondiral function and reducing oxidative stress, and *Plxna2* and *Gja1*, which are implicated in vascular integrity^[14]^. Genes related to modulation of VEGF signalling like *Adamts1, Adamts6,* and *Sema6d^[15]^* were upregulated as well. Overall, these data suggest that in addition to reducing vasculature permeability^[12a]^, the iECM also helps to maintain endothelial integrity and modulate VEGF-induced angiogensis. When these genes were analyzed via GO (Figure 2C), enriched pathways included vasculature and tube development, and actin filament-based processes. Saline-treated animals also exhibited some angiogenic markers such as *Foxp1* and *Notch1* which is an expected result post-MI^[16]^. However, saline also saw upregulation of genes such as *Plxnd1*, *Ddx5* and *Fabp4* which are closely related to endothelial cell injury, scar formation and pro-inflammatory M1-like macrophage infiltration*^[17]^*. In addition to angiogenic and DNA damage responses in saline-treated animals, GO analysis revealed terms implicating endothelial cell migration and anchoring junctions (Figure 2C). The upregulated genes (*Plxnd1, Ddx5,* and *Fabp4*) and GO terms both suggest negative post-injury vascular remodeling, characterized by neointima formation and pro-inflammatory chemoattractant signals, which may exacerbate edema immediately post-MI^[18]^. This gene expression profile contrasts starkly with the pro-reparative signals and positive vascular maintenance seen in the endothelial cells of iECM-treated animals at day 1 post-MI.

At day 3, a similar endothelial cell behavior can be seen. The endothelial cell population as a whole shows a generally angiogenic response via upregulation of *Pecam1^[19]^*, *Tek^[16b]^*and *Sema6a^[15]^*. In addition, the upregulation of *Dock6* and *Dock9* suggest cytoskeletal reorganization of endothelial cells in support of vasculature development while *Adgrl4^[20]^* and *Adgrf5^[21]^* suggest vascular barrier formation and endothelial cell adhesion. These findings are further supported by the GO enrichment analysis (Figure 2C) which found enrichment for angiogenesis, vasculature development and cell migration. Saline treated endothelial cells at day 3 showed a similar remodeling behavior as day 1 with upregulation of *Ddx5*, *Col3a1, Actb*, *Vim, Itgb1*, and *Adamts6/9.* Gene ontology enrichment (Figure 2C) showed signs of cell morphogenesis and DNA damage response. Overall, the saline treated endothelial cells showed that the lingering angiogenic behavior seen at day 1 was beginning to diminish and a more remodeling/fibrotic response was beginning to appear.

At day 7, when leaky vasculature is significantly reduced and iECM has already degraded^[12a]^, scRNAseq reveals the prolonged pro-reparative effects of the iECM. Genes related to angiogenesis like *Notch1*, *Rasip1,* and *Srsf2* and reduced apoptosis like *Notch4* are upregulated with iECM treatment ^[16b, 16c, 22]^. In addition, genes related to the basement membrane like *Ccnl2* and endothelial cell adhesion like *Arghef15* are upregulated *^[23]^*. In saline-treated animals, *Fabp4* remained upregulated as observed on day 1, indicating a pro-fibrotic phenotypic shift through endothelial cell to mesenchymal transition towards myofibroblasts^[24]^. *B2m*, a biomarker associated with heart failure, was also elevated^[25]^. Saline treatment also yielded upregulation of *Aqp1*, which is related to myocardial edema^[26]^, confirming that lasting effects of MI-induced leaky vasculature are present without iECM treatment. GO showed enrichment for vasculature development and blood vessel morphogenesis for iECM treatment, which strongly contrasts with saline-treated animals, whose endothelial cells at day 7 were enriched for increased leukocyte cellular adhesion and sustained anchoring junction dysfunction (Figure 2C). Finally, we compared iECM effects across time by performing pseudobulk analyses between day 1 and day 3 iECM samples (Figure 2D) with day 1 and day 7 iECM samples (Figure 2E). With both comparisions, iECM elicited markers of angiogenesis (*Hif1a, Adamts1, Actg1, Adamts9*) at day 1 post infusion, and promoted antioxidative reactions at both days 3 and day 7 post infusion (*Zbtb16, Myrip).* These differentially expressed genes are displayed in Table S3 (Day 1 vs. Day 3) and Table S4 (Day 1 vs. Day 7).

Similarly, we found responses with the lymphatic endothelium (subsetted via *Prox1+/Flt4+ expression*) post subclustering of various populations at day 1, day 3 and at day 7 (Figure S5A-C). The lymphatic subcluster markers are outlined in Table S1 and the DEGs outlined in Table S5. At day 1, lymphatic endothelial cell development was seen in both treatments such as upregulation of *Sulf1*, *Myct1*, *Hif1a, Glis3,* and *Angpt2* within the iECM*^[27]^* and *Ccl21*, *Tspan18*, *Lyn, Mtss1, Tspan18,* and *Notch1* in saline treatment*^[28]^*. Overall, there were few DEGs across treatment and no enriched GO pathways (Fig. S5C). At day 3, few DEGs were found suggesting a lack of lympahtic endothelial cell development, mirroring the day 1 behavior. By day 7, significant differences in the lymphatic endothelial cells were seen. Within iECM treatment, genes for lymphatic development were found such as *Fn1*, *Nrp2* and *Tspan18^[29]^*. Moreover, genes tied to cardiac remodeling and immunosuppression like *Tbx1*, *Plcg1*, and *Ahr* were upregulated as well*^[30]^*. Via GO, pathways for vasculature development, basement membrane and wound healing were enriched. Within saline treatment, like endothelial cells, *Aqp1* was upregulated showing the lasting effects of edema. While saline treatment also shows some pro-reparative gene expression via *Efnb2^[31]^,* which is part of the normal reparative reaction post-MI, there was also upregulation of pro-apoptotic *Igfbp5^[32]^*. This was ultimately reflected in enriched GO terms such as decreased regulation of cellular differentiation and response to oxygen compounds, which may indicate oxidative stress (Figure S5C). We then evaluated how the iECM promotes lymphatic endothelial cell development over time by comparing the iECM response at Day 3 and Day 7 relative to Day 1 (Figure S5D-E). At day 1, we found a conserved angiogeneic *(Hif1a*) and developmental response *(Adamts9*). At both days 3 and day 7, genes involved in further angiogenic and survival responses (*Etl4*, *Prkag2*) alongside ECM interactions (*Tmtc1*, *Eln*) were found. These differentially expressed genes are displayed in Tables S6 (Day 1 vs. Day 3) and S7 (Day 1 vs. Day 7).

### 2.4. Mural Cells Exhibit Vasculature Developmental Gene Phenotypes due to iECM

Closely tied to the repair of the vasculature are smooth muscle cells and pericytes, which have been shown to proliferate and develop in response to ECM administration^[33]^. Due to the similarity in gene expression and flow association within the vasculature surrounding endothelial cells, pericytes and vascular smooth muscle cells were subset together using *Myh11+/Tagln+/Acta2+,* clustered (Figure S6A-C, S7A-C) and quantified relative to treatment condition (Figure S4B), and analyzed together as mural cells with their UMAPs (Figure S6A), DEGs (Figure S6B) and enriched GO terms (Figure S6C) displayed. Mural cell DEGs across time are outlined in Table S8.

At day 1, mural cells with iECM treatment showed gene expression related to angiogenesis through *Robo1*, *Snd1* and *Ext1* upregulation*^[34]^*, and cardioprotection via *Osmr*^[35]^. Pathway enrichment supported these findings showing cell motility and GAG metabolic process. This further supports that idea that the iECM is able to induce angiogenesis and impact the vasculature in a pro-reparative way acutely post-MI similar to its effect on endothelial cells. Saline treated mural cells also exhibited upregulation of *Ddx5* as well as *Septin4* and *Cd63* which are tied to reduced proliferation and fibrosis *^[36]^*.

At day 3, mural cells with iECM treatment did not show much evidence of development and did not enrich for any GO pathways. With saline treatment, mural cells showed signs of reduced vasculature development via *Ddx5* upregulation in addition to significant ECM remodeling genes like *Col4a1, Fbn1*, *Actb, Col3a1* and *Col1a2*. Overall, this suggested that while iECM treatment did not significantly impact mural cell development at day 3, the saline treatment exhibited a significantly higher remodeling and fibrotic phenotype.

At day 7, mural cells continued to exhibit upregulation of genes tied to migration, proliferation and angiogenesis such as *Col6a1 Flna, Rgs6* and *Pdgfrb^[37]^*. In contrast, mural cells of saline treated animals exhibited a mixed expression profile, including upregulation of both *Igf1^[38]^*, associated with cardioprotection, and *Tmod3^[39]^*, associated with reduced angiogenesis. In iECM treated animals, the DEGs of mural cells at day 7 were enriched for basement membrane matrix and platelet derived growth factor binding, implicating the continuation of angiogenesis and vasculature development. We then evaluated differentially expressed genes for iECM treated samples between Day 1 and Day 3 post infusion, and Day 1 and Day 7 post infusion. At day 1, we found conserved markers involved in development (*Adamts9*). Both days 3 and 7 also have conserved markers for angiogenesis and proliferation regulation (*Elt4, Rora*). These differentially expressed genes are displayed in Table S9 (Day 1 vs. Day 3) and Table S10 (Day 1 vs. Day 7). Overall, these findings indicate that the iECM elicits a proliferative and angiogenic response within the myocardium shortly after MI, which is sustained through day 7.

### 2.4. iECM Directly Affects Immune Cell Types at Day 1

Decellularized ECM biomaterials such as iECM have unique properties that elicit beneficial effects post-injury via immunomodulation^[11a, 11b, 11e, 12, 40]^. Early post-MI, monocytes are among the first immune cells to infiltrate and differentiate into macrophages in addition to neutrophils. To understand iECM’s effect on the immune cells immediately present within the infarct, monocytes (*Cxcr2+/Csf3r+)* and antigen presenting cells (*Cd74+/Ciita+)* were subsetted, reclustered, and visualized in UMAP space **(Figure 3A-B)** with subcluster markers in Table S1, and iECM vs. saline analysis displayed in Table S11. The volcano plots are displayed (Figure 3C-D) with enriched GO terms (Figure 3E-F). These phenotypes were largely depleted day 1 post-MI, consistent with typical activation and differentiation patterns of monocytes^[4]^.

**Figure 3.**
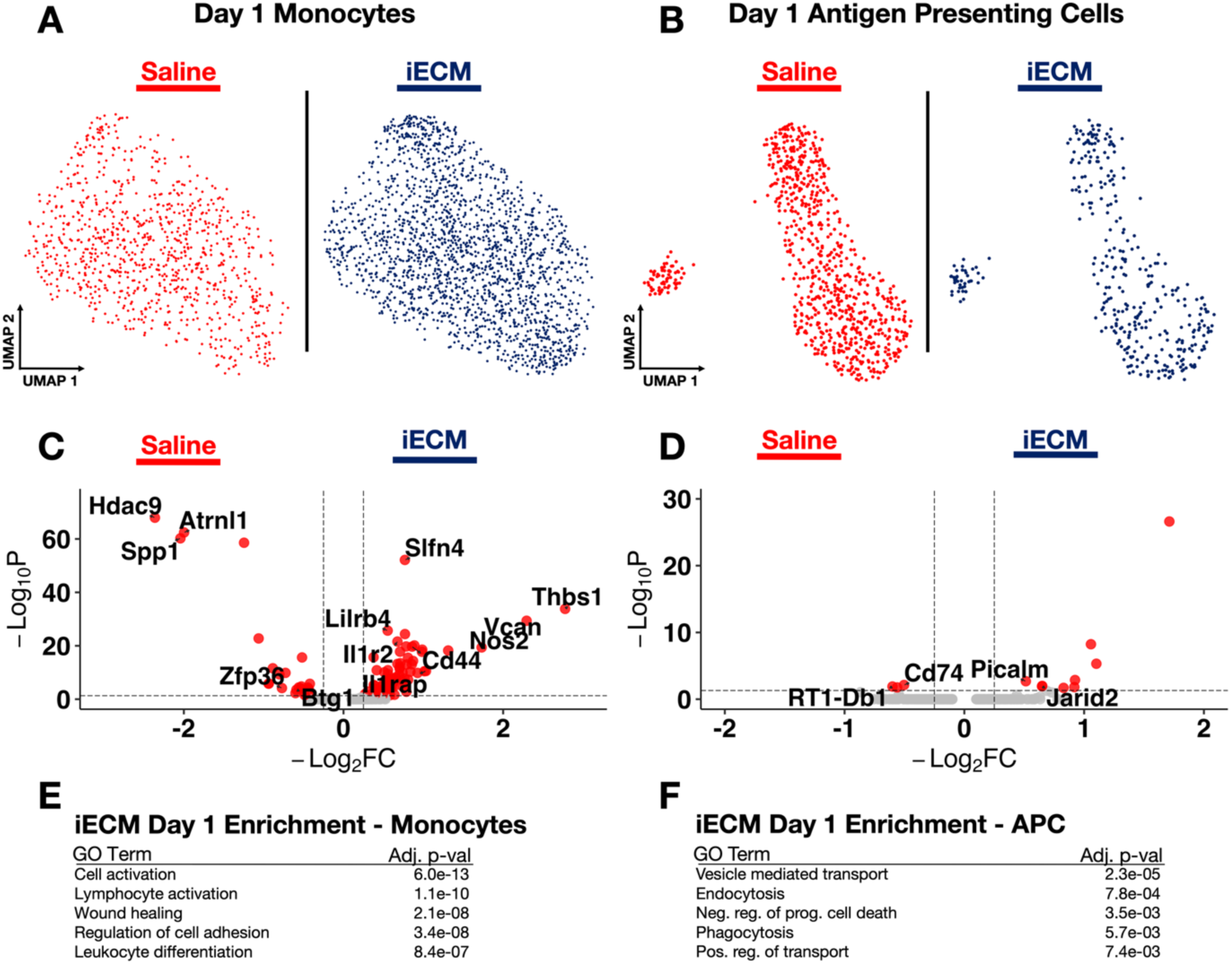
iECM impacts early immune cell phenotypes 1-day post-MI. UMAPs of iECM and saline for A) monocytes and B) antigen presenting cells. Volcano plots of DEGs for C) monocytes and D) antigen presenting cells. Pathways enriched via GO for E) monocytes and F) antigen presenting cells.

Early monocytes in the iECM-treated infarcts strongly expressed genes encouraging innate immune cell regulation (*Thbs1, Cd44) ^[41]^,* anti-inflammatory macrophage polarization (*Vcan, Nos2, Slfn4) ^[42]^,* the blockade of the inflammatory interleukin-1b pathway (*Il1rap, Ilr2)^[43]^,* and the limitation of T cell infiltration (*Lilrb4)^[44]^.* In stark contrast, early myeloid cells in saline controls expressed genetic regimes associated with pro-inflammatory macrophage polarization (*Hdac9, Spp1) ^[45]^*, ischemic heart failure (*Atrnl1)^[46]^,* B-cell recruitment, and upregulated apoptosis (*Btg1)^[47]^*. GO revealed enrichment of terms relating to lymphocyte activation and differentiation, wound healing, and cell activation in iECM-treated early non-macrophage myeloid infiltrators. These data suggest the infiltrating monocytes and neutrophils play a role in modulating the immune response.

The early anti-inflammatory response with iECM treatment may be attributed to the protective signaling from antigen presenting cells (APCs) captured at 1 day post-MI, in addition to the early monocytes mentioned previously. APCs from iECM-treated infarcts exhibited differentially expressed genes involved in anti-inflammatory leukocyte polarization (*Slfn4)*, and attentuation of antigen presentation and immune cell function (*Htr7, Jarid2, Pitpnc1, Atp6v0a1*). In contrast, saline-treated APCs showed strong differential expression of activated major histocompatibility complex (MHC) class II genes *(RT1-Db1, Cd74,* and *Hdac9)^[48]^*. iECM bolstered the pro-reparative response in APCs and macrophages at day 1 post-infusion, encouraging a sustained temporal shift in the anti-inflammatory polarization of leukocytes. This early immunomodulation may contribute to the beneficial differences observed in other myocardial phenotypes post-MI.

### 2.5. The Temporally Distinct Macrophage Response Post-MI Is Expedited through iECM Treatment

Given that iECM encourages the earliest myeloid cell infiltrators to adopt pro-reparative differentiation schemes and anti-inflammatory signaling, we subsetted and refined macrophage clusters (*Mrc1+/Csf1r+)* captured via snRNAseq. Cells at each timepoint were integrated and projected onto UMAP space **(Figure 4A)**. Subclusters were identified (Figure S8), and we measured proportions with respect to treatment (Figure S4C), with marker genes displayed in Table S1. Here, we also subsetted with respect to origin, as macrophages can be monocyte-derived (*Cd74+/Ccr2+*) or resident cardiac macrophages (*Mrc1+)*. For downstream comparisons, iECM vs. saline macrophages demonstrated transcriptional activation of anti-inflammatory gene expression as early as 1 day post-infusion, and sustained up to 7 days (Figure 4B).

**Figure 4.**
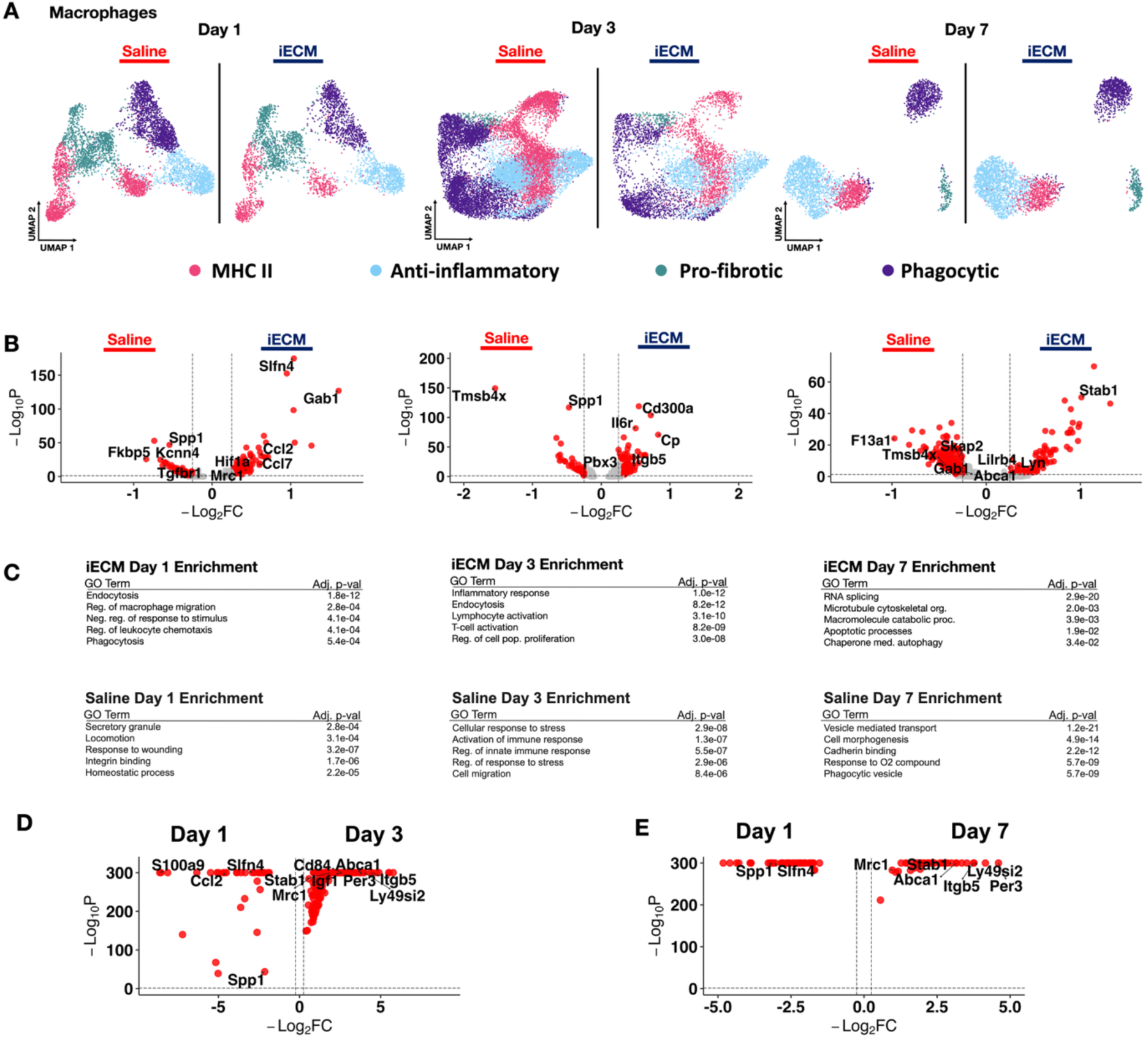
Macrophages demonstrate acute anti-inflammatory responses with iECM treatment. A) UMAP of macrophages in iECM and saline treated samples at 1-, 3-, and 7-days post infusion. B) Differentially expressed genes of iECM and saline macrophages. C) All iECM specific differentially expressed genes were subjected to GO enrichment. D) Differentially expressed genes in macrophages with iECM treatment at day 1 versus day 3 post infusion. E) Differentially expressed genes in macrophages of iECM treatment at day 1 versus day 7 post infusion.

At day 1 post-infusion, we measured genes corroborating a largely M2-like pro-reparative phenotype (*Gab1, Ccl2, Mrc1, Mertk, Hif1a and Slfn4)^[48c, 49]^,* although some pro-inflammatory phenotypes were also observed (*Ccl7)^[50]^.* In comparison, saline macrophages exhibited high pro-inflammatory markers that contribute to pro-fibrotic M1-like macrophage polarization *(Spp1, Tgfbr1, Kcnn4, Fkbp5)* at day 1 post infusion*^[45b, 51]^*. At day 3, iECM continued to encourage pro-reparative polarization (Figure 4B), with macrophages expressing key M2-like DEGs *(Stab1, Slc9a9).* Macrophages from saline-treated animals had enhanced expression of fibrosis and phagocytosis markers *(Spp1, Vim, Col3a1).* The anti-inflammatory polarization in iECM-treated macrophages is sustained until day 7 with *Stab1, Lyn,* and *Lilrb4.* Saline-treated macrophages exhibit a mixed response, expressing markers related to dendritic cell activation with *Tmsb4x*, while simultaneously displaying M2-like polarization with the moderate expression of *F13a1, Skap2,* and *Gab1^[52]^.* These responses were validated through GO, which emphasized the regulation of leukocyte chemotaxis and macrophage migration at day 1, lymphocyte activation and cellular proliferation regulation at day 3, followed by apoptotic processes and chaperone-mediated autophagy at day 7 post-infusion with iECM treatment (Figure 4C). Saline-treated macrophages were enriched with terms such as locomotion, responses to wounding, and secretory granules at day 1, followed by enriched responses to O_2_ compounds and phagocytic vesicles (Figure 4C). All differentially expressed genes here are outlined in Table S12.

Given that ECM biomaterials are known to elicit anti-inflammatory responses^[11e, 53]^, we then measured the specific anti-inflammatory macrophage response between iECM and saline samples. At day 1 post infusion, the iECM elicited significant anti-inflammatory responses, demonstrated by *Mrc1, Cd163,* and *Il4r* expression (Figure S9A). This response is sustained even at day 3 post infusion as demonstrated with expression of *Cp, St3gal5,* and *Cd300a* (Figure S9B). At day 7 post infusion, the iECM still elicited high *Stab1* expression and downregulation of *Il6r,* demonstrating continuation of the anti-inflammatory response (Figure S9C). All differentially expressed genes are displayed in Table S13. Comparisons across time for iECM samples were also done. At day 1, there were genes involved in the neutrophil response (*Csf3r, S100a8, S100a9*) and general inflammation (*Cxcr2, Ccl2*) (Figure 4D-E). At both day 3 and day 7, there are markers for anti-inflammatory macrophages *(Mrc1, Stab1*) (Figure 4D-E). All differentially expressed genes over time are displayed in Table S14 and S15 for comparisons between Days 1 and 3 post infusion and Days 1 and 7 post infusion, respectively.

### 2.6. iECM Elicits Fibroblast Activation

The immune response is also known to directly mediate other cellular responses to elicit cardiac repair and remodeling^[1b, 54]^. Previously, we demonstrated that a decellularized myocardial ECM hydrogel elicits fibroblast activation^[11e, 53]^. Fibroblasts were examined similarly by subsetting them via *Gsn+/Dsn+* expression and subclustering them **(Figure 5A)**. The subcluster populations were identified (Figure S10), the relative subcluster populations were found (Figure S4D), and the marker genes are displayed in Table S1. We then ran DEG analysis (Figure 5B), and evaluated the DEGs via GO enrichment (Figure 5C). All fibroblast DEGs are outlined in Table S16.

**Figure 5.**
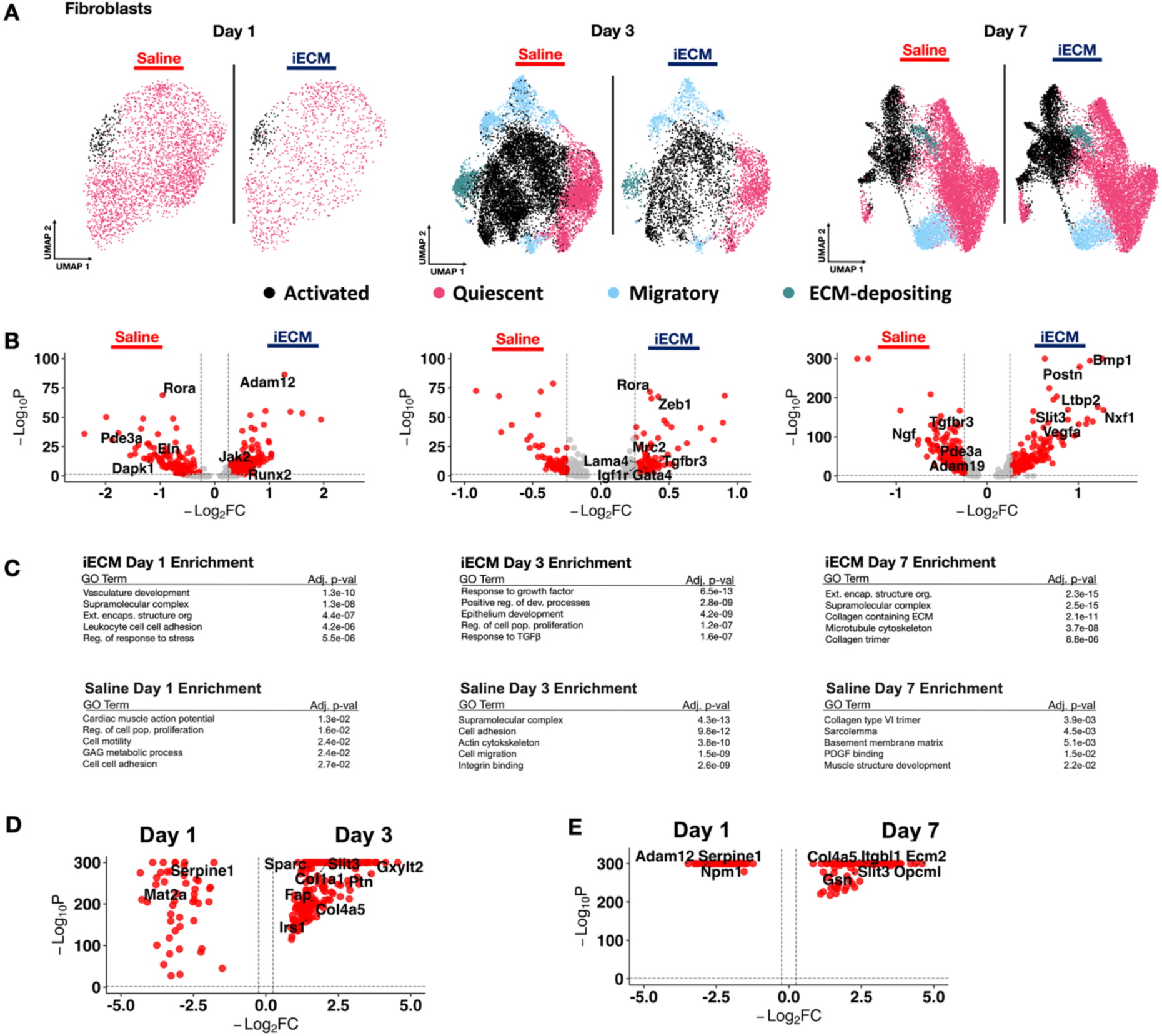
iECM promotes fibroblast activation. A) UMAP of fibroblasts in iECM and saline treated samples split across days 1, 3 and 7 post infusion. B) Differentially expressed genes of iECM and saline fibroblasts at days 1, 3 and 7 post-infusion. C) All iECM specific differentially expressed genes in the fibroblast subset were subjected to GO enrichment. D) Differentially expressed genes of fibroblasts in iECM samples at day 1 versus day 3 post infusion. E) Differentially expressed genes fibroblasts in iECM samples at day 1 versus day 7 post infusion.

As a major cell type within the myocardium, the fibroblasts at day 1 exhibited gene expression profiles indicative of immune signaling activity via upregulation of *Pkm^[55]^*. Additionally, genes such as *Runx2* and *Adam12*, implicated in mitigating cardiac hypertrophy and remodeling*^[56]^*, were upregulated. Playing a role in vasoregulation post-MI*^[57]^*, *Kcnq5* was also found to be upregulated. When run through GO analysis, pathways related to leukocyte cell adhesion, regulation of stress response, and enrichment of ECM structurural organization were revealed, supporting the notion that iECM treatment promotes an immunoregulatory, pro-reparative fibroblast phenotype. In contrast, saline treated fibroblasts showed enrichment of genes associated with cardiac remodeling and hypertrophy such as *Rora*, *Eln, Dapk1*, and *Pde3a ^[58]^.* This was further supported by GO analysis, which revealed enrichment of pathways for cardiac muscle action potential, cell motility, and cell-cell adhesion, suggesting maladaptive cardiac morphological changes (Figure 5C).

At day 3, iECM treated fibroblasts showed upregulation of *Tgfbr3, Rora, Zeb1, Mrc2, Lama4, Igf1r, and Gata4.* At this timepoint, these fibroblasts exhibit migratory markers (*Zeb1*), ECM remodeling markers (*Mrc2, Gata4, Lama4*), and pro-reparative remodeling (*Tgfbr3, Rora, Igf1r)*. This is further demonstrated with GO analysis, which showed response to growth factor pathways, such as TGF-β.

At day 7, fibroblasts in iECM treated infarcts showed upregulation of *Vegfa* suggesting a prolonged impact on promoting angiogenesis. However, iECM treated animals also showed upregulation of genes tied to fibrosis such as *Slit3* and *Bmp1^[59]^.* These DEGs are corroborated by GO enrichment for collagen containing ECM and collagen trimer. The saline treated animals exhibited a mix of pro-fibrotic markers like *Pde3a* and pro-reparative markers like *Tgfbr3,* which matched the GO enrichment for basement membrane matrix and muscle structure development (Figure 5C). As fibroblast remodeling is a key facet in MI pathophysiology^[11e, 60]^, we then evaluated the activated fibroblast response (Figure S11). At day 1 post infusion, we found that the iECM elicited less fibroblast activation via downregulated *Postn*, *Col1a1,* and *Col3a1* expression, all of which are markers for further fibroblast activation and ECM remodeling^[8a]^ (Figure S11A). Similarly, at day 3 post infusion, we also see markers of activated fibroblasts with *Tnc and Fn1* in saline treatment (Figure S11B). However, at day 7 post infusion, we measured higher fibroblast activation with iECM treatment, as represented with *Col1a1, Tnc, Bmp1,* and *Ltbp2* upregulation (Figure S11C). All differentially expressed genes are displayed in Table S17.

We also evaluated differentially expressed genes for iECM treated samples between Day 1 and Day 3 post infusion alongside comparing Day 1 and 7 post infusion. We found conserved markers of cell adhesion (*Ptges3, Pkm, Myob)* at day 1, higher fibroblast activation at day 3 (*Sparc, Gap, Col1a1)*, and markers of ECM organization (*Col3a1, Col4a3, Fbln1, Col14a1)* at day 7 (Figure 5D-E). All differentially expressed genes over time are displayed in Table S18 for comparing Day 1 and Day 3 post infusion and Table S19 for comparing Day 1 and 7 post infusion. Overall, these results suggest that the iECM impacted the immune response and repair response within fibroblasts early in the acute phase post-MI. While some of the pro-reparative effects are preserved, the overall injury and downstream fibrotic response are not ablated.

### 2.8. iECM Elicits Myocardial Salvage and Developmental Genes in Cardiomyocytes

The iECM has also been shown via histology to have decreased cardiomyocyte apoptosis within the infarct border zone^[12a]^. To explore this phenomenon at the cellular level, we analyzed the snRNAseq dataset by isolating, reclustering (Figure S12) and identifying relative proportions (Figure S4E), and splitting cardiomyocytes (*Rbm20+/Ryr2+*) based on iECM and saline treatment, followed by subjecting the data set to DEG and GO analysis (**Figure 6**). The DEGs identified in cardiomyocytes from both groups across the acute timeframe are displayed in Table S20.

**Figure 6.**
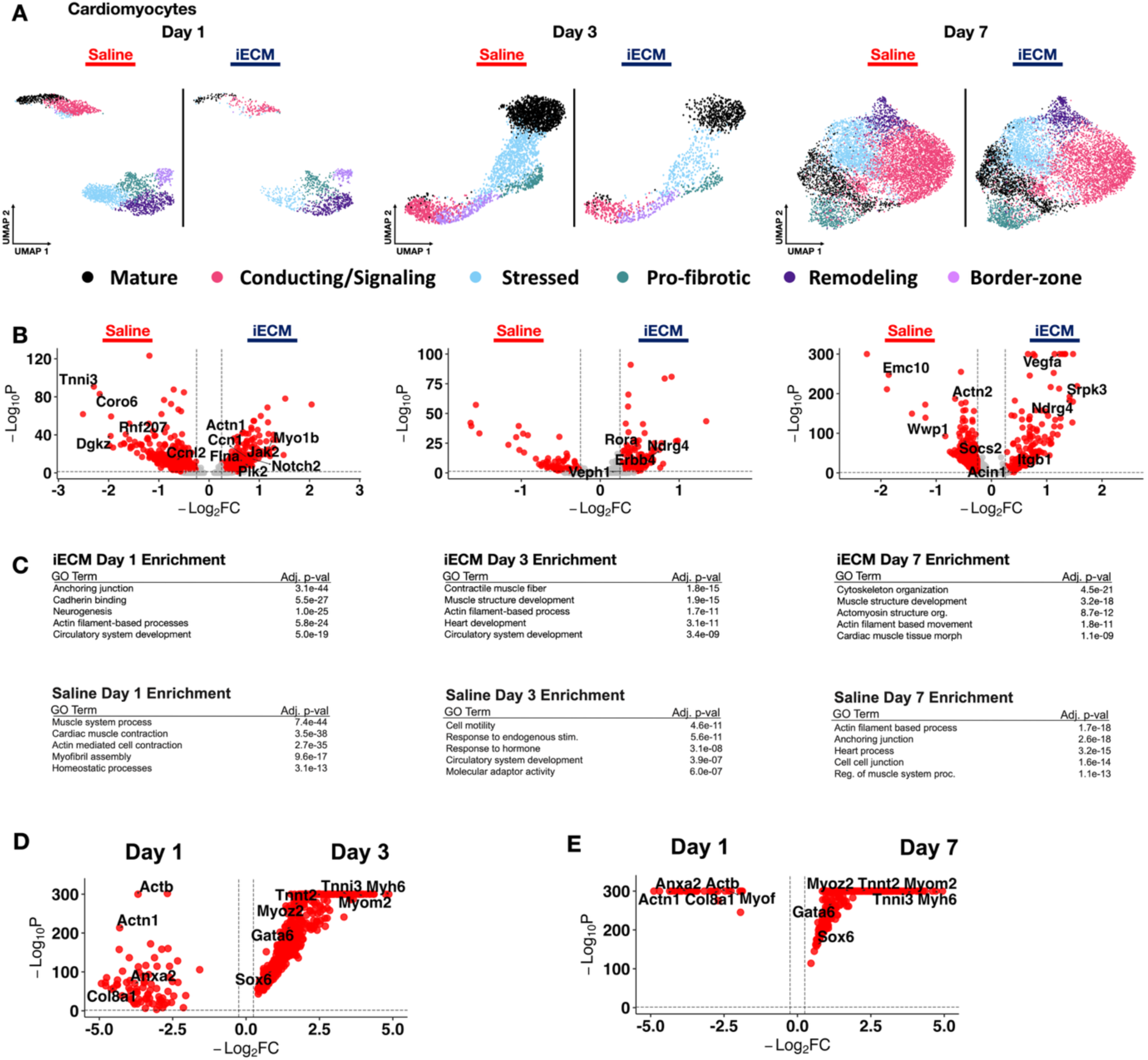
iECM promotes myocardial salvage. **A)** UMAP of cardiomyocytes in iECM and saline treated samples across days 1, 3 and 7 post infusion. **B)** Differentially expressed genes in cardiomyocytes from iECM versus saline samples at days 1, 3 and 7 post-infusion. **C)** All iECM specific differentially expressed genes in the cardiomyocyte subset were subjected to GO enrichment. D) Differentially expressed genes in cardiomyocytes with iECM at day 1 versus day 3 post infusion. E) Differentially expressed genes in cardiomyocytes with iECM at day 1 versus day 7 post infusion.

At day 1, a variety of genes linked to reduction of apoptosis, reduction of fibrosis and cardiomyocyte proliferation were upregulated with iECM treatment. In particular, *Notch2, Hsp90b1, Myo1b*, and *Jak2* which are tied to the reduction of apoptosis post-MI*^[61]^*, were upregualted with iECM treatment. Genes tied to cardiomyocyte proliferation like *Ccn1* and *Ccnl1^[62]^* and genes linked to reduction in fibrosis, *Plk2* and *Flna^[63]^*, were also upregulated. In contrast, saline treated animals showed upregulation of genes like *Rnf207*, which is linked to cardiac hypertrophy*^[64]^*, and *Dgkz*, which is a marker for coronary artery disease progression*^[65]^*. However, saline treated animals also showed upregulation of *Myom2^[66]^*, which is crucial for heart function. Via GO, day 1 cardiomyocytes treated with iECM were enriched for anchoring junction, neurogensis, actin filament-based processes, and circulatory system development, suggesting the preservation of cardiac function post-MI. We previously demonstrated that the iECM indeed improves cardiac function as early as day 1 post-MI^[12a]^.

At day 3, the cardiomyocyte response with iECM treatment was maintained with upregulation of *Fgf1,* which improves angiogenesis within the myocardium^[67]^, *Rora*, which is implicated in reducing oxidative stress^[68]^, and *Ndgr4*, which is linked to the Akt signaling pathway^[69]^. This was further supported by GO analysis showing enrichment for heart development and circulatory system development suggesting that iECM showed prolonged cardiomyocyte development. Saline treated cardiomyocytes at day 3 also showed signs of cardiomyocyte development and survival via *Stat3^[70]^*, *Zeb1^[71]^,* and *Foxp1^[16a]^* upregulation. The particular GO enrichment supported these findings with circulatory system development.

At day 7, cardiomyocytes from iECM treated animals showed DEGs related to cardioprotection via upregulation of *Srpk3* and *Acin1*, which are both tied to oxidase-mediated cardioprotection^[72]^ and *Ndgr4*, similar to day 3^[69]^. These cells also showed upregulation of *Vegfa^[73]^* suggesting preserved cardiac angiogensis as late as day 7 post-MI and the appearance of *Itgb1*, which has been shown to increase stem cell survival and improve cardiac function post-MI^[74]^. GO enrichment further supported these findings with pathways for muscle structure development, actomyosin structure organization, and cardiac muscle tissue morphogenesis. In contrast, saline treated cardiomyocytes showed upregulation of *Wwp1*, which is linked to inflammation^[75]^ and *Actn2*, which is linked to heart failure^[76]^. This also suggests that iECM is able to preserve the cardioprotective effects seen early on at day 1 and day 3. We also measured differentially expressed genes for iECM treated cardiomyocytes (Figure 6D-E). We found conserved adhesion markers (*Anxa2, Adam12, Col8a1*) and actin-based processes (*Actn1, Actb*) at day 1. Both day 3 and day 7 exhibited markers of actin-based processes (*Myom2, Myoz2*), healthy cardiomyocytes (*Myh6, Tnnt2*), and muscle development (*Gata6, Sox6*) (Figure 6D-E). All differentially expressed genes are displayed in Table S21 (Day 1 vs. Day 3) and Table S22 (Day 1 vs. Day 7). Overall, the pro-reparative effects of the iECM were found to enhance cardioprotection, promote development, and actin-based processes in the cardiomyocytes. While these markers are already present as early as 1 day post-infusion, these changes are more pronounced over time, particularly at day 7 post-infusion.

### 2.9. Neurogenesis and Neuronal Cell Development Are Exhibited with iECM Administration

Previous studies with our ECM hydrogel suggested a therapeutic effect on the neuronal cells present in the infarcted heart, particularly in the subacute and chronic phases of MI^[11e]^. In a similar fashion, we thus measured iECM therapeutic effects on neuronal cells, crucial for conduction and action potential transduction during myocardial contraction. We subsetted neuronal cells via *Ncam*+/*Cadm2+* with marker genes displayed in Table S1. We mapped and analyzed them via DEG and GO analysis similarly (Figure S13A-C). All differentially expressed genes for neuronal cells across time are outlined in Table S23.

At day 1 and day 3, the neuronal response was weak with few DEGs and no GO enrichment. This is likely due to the delay in cardiomyocyte development which was predominantly seen via DEG analysis at day 7. Fittingly, day 7 neuronal cells showed markers for proliferation and neurogenesis through upregulation of *Npdc1*, *Agrn,* and *Dst^[77]^*. Furthermore, GO enrichment revealed basement membrane and structural molecule activity suggesting the development and stabilization of neuronal cells within the myocardium. In contrast, neuronal cells in saline treated infarcts also showed signs of neuronal development via upregulation of *Ntng1* and *Lrrtm3^[78]^*, however, there was also high expression of *Pmp22^[79]^*, which is a marker of congestive heart failure post-MI. Similar to the full ECM hydrogel, this shows that the infusible ECM can elicit markers of neurogenesis. Lastly, iECM treated neuronal cells at days 1 and 3 post infusion and days 1 and 7 post infusion were subjected to comparison. For both comparisons, day 1 exhibited markers that lessened development and heightened inflammation (*Il6st*), while days 3 and 7 exhibited markers of neuron development (*Ntng1, Nrxn1, Grid2*) (Figure S13D-E) with all genes displayed in Tables S24 (Day 1 vs. Day 3) and S25 (Day 1 vs. Day 7).

### 2.10. iECM Induces Pro-Reparative Effects Spatially Within the Infarct Across Time

Finally, given the iECM is distributed throughout the MI vasculature and is known to localize to the infarct^[12a]^, we evaluated therapeutic activity through spatial transcriptomics by comparing the infarct zones of an iECM treated section to saline treatment. Infarct zones were determined through unsupervised clustering, with infarct specific clusters containing low *Myh6* and *Tnnt2* expression (Figure S14), allowing for us to differentiate spots that are infarct specific in both iECM and saline treated samples **(Figure 7A)**. After integrating spatial samples with both conditions, we plotted the DEGs between iECM and saline treatment across time.

**Figure 7.**
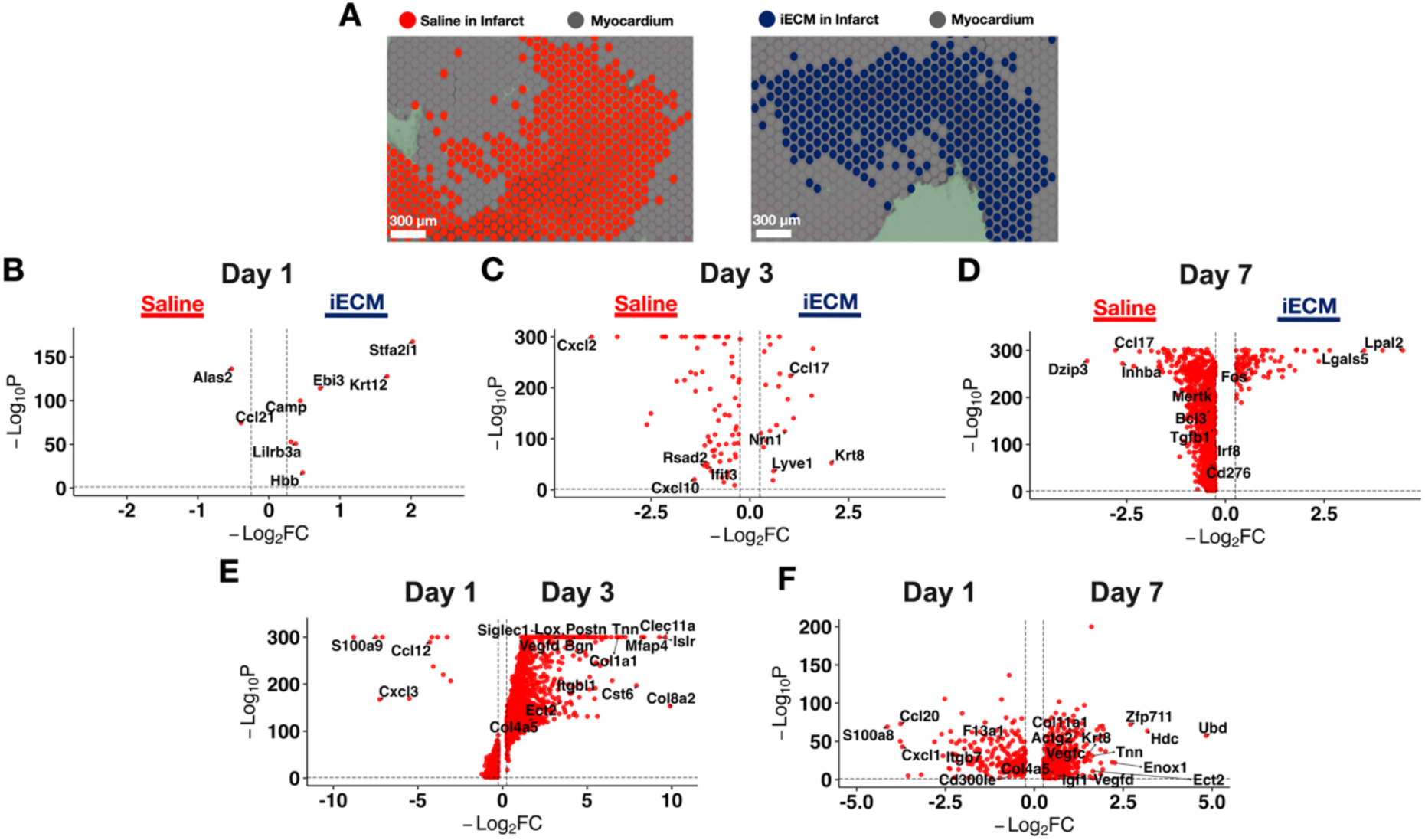
iECM exhibits subtle spatial differences with respect to saline across time. A) 10X Visium sections indicate the spatial difference between iECM treated infarcts (red) to saline treated infarcts (cyan). B) Volcano plot displays differences between iECM and saline treated spatial transcriptomic sections at 1 day post infusion. C) Volcano plot displays differences between iECM and saline treated spatial transcriptomic sections at 3 days post infusion. D) Volcano plot displays differences between iECM and saline treated spatial transcriptomic sections at 7 days post infusion. E) Volcano plot displays differences between iECM at 1-day vs. 3-day post infusion. F) Volcano plot displays differences between iECM at 1-day vs. 7-day post infusion.

At 1 day post-infusion (Figure 7B), iECM exhibited high expression of *Camp*, *Lilrb3a, Krt12, and Stfa2l1*. Saline treated hearts expressed high *Ccl21*, a chemokine that plays a crucial role in recruiting immune cells, such as inflammatory T-cells^[80]^ (Figure 7B). Here, through spatial transcriptomics, we thus see that the iECM mitigates the pathological immune response, as reflected through the anti-inflammatory macrophage response, innate immune cell regulation, and blockage of pathological T-cell activation in snRNAseq.

At 3 days post-infusion (Figure 7C), iECM exhibited genes such as *Krt8*, a marker of TNF induced protection^[81]^. In contrast, the saline differentially expressed genes displayed markers involved in the pro-inflammatory innate immune response (*Ifit2, Ifit3, Isg15, Rsad2)*^[82]^. At 7 days post-infusion (Figure 7C), iECM demonstrated elevated *Ccl20, Lgals5, Lpal2,* and *Fos,* all markers representing regulation of the immune response and overall cell proliferation^[83]^. In addition, GO revealed regulation of programmed cell death, and regulation of the stress response (Figure S15B). Here, spatial transcriptomics corroborates the sustained anti-inflammatory response found with iECM administration, reduced apoptosis found in cardiomyocytes and the endothelium, and response to stressors, such as O_2_ compounds, found in the snRNAseq data. Saline exhibited genes with increasing pro-inflammation (*Mertk, Irf8*)^[84]^, and markers of negative remodeling in MI (*Dzip3, Inhba)^[85]^.* The snRNAseq macrophage response in saline treated animals at day 7 largely validates the spatial data, with further markers of pathological remodeling measured in saline treated endothelium, fibroblasts, mural cells, cardiomyocytes, and neuronal cells. All differentially expressed genes comparing the iECM and saline infarct zones at each timepoint are outlined in Table S26.

We then evaluated the iECM spatial differences over time by comparing iECM treated infarcts at days 1 and 3 post infusion (Figure 7D) and a similar comparison between days 1 and 7 post infusion (Figure 7E). Day 1 post infusion iECM infarcts elicited significant pro-inflammatory markers, such as *S100a8, Cxcl1, Ccl20*, and *Cd300le* (Figures 7D-E), which was conserved when comparing Days 1 and 3 alongside Days 1 and 7 in tandem. However, at day 3 post infusion, we measured higher anti-inflammatory immune specific responses, particularly with *F13a1* and *Siglec1* expression, and fibroblast and ECM specific activity (*Postn, Lox, Bgn1, Col1a1*). When comparing days 1 and 7 post infusion, day 1 elicited anti-inflammatory immune specific responses, particularly with *Arg1, F13a1,* and *Gata3* expression. In comparison, at day 7 post infusion, there were markers of tube development via *Vegfc, Vegfd,* and *Slit1* expression, cardioprotection via *Igf1*, and fibroblast activation and ECM composition via *Col11a1, Col4a4, Col4a5*, and *Hmcn1*. All significant genes across time are displayed in Tables S27 (Day 1 vs. Day 3) and S28 (Day 1 vs. Day 7).

Finally, as a metric of further validating the differentially expressed genes present through snRNAseq, we mapped out the iECM specific genes associated with each timepoint directly onto the infarct of iECM and saline treated spatial samples. We found that the iECM DEGs from endothelial cells, fibroblasts, and cardiomyocytes were significantly higher in iECM treated spatial samples at days 1 (Figure S16A) and 3 (Figure S16B) compared to saline treated spatial samples. At day 7, in addition to fibroblasts and cardiomyocytes, mural cell iECM DEGs scored significantly higher in iECM spatial samples than saline (Figure S16C).

## 3. Discussion

Here, we show the pro-reparative effects across time of an intravascularly administered decellularized biomaterial. We were able to validate previous findings that the iECM elicits pro-repair programs such as immunomodulation, vasculature development, and myocardial salvage in acute MI^[12a]^. While we acknowledge that the GO analyses should be interpreted with caution, these GO terms are consistent with previously published findings for our infusble ECM^[12a]^, and with our full ECM hydrogel^[9a, 9b, 53]^. We also note newer findings with the iECM, such as macrophage polarization, fibroblast activation, induction of a myocardial developmental transcriptomic program, neurogenic responses, and effects on lymphatic development. Due to intravascular administration of the iECM and its localization in small micron scale areas, we could not measure spatial heterogeneity within the infarct as with ECM hydrogels or other biomaterials^[11b, 11d]^. Regardless, iECM upregulated genes involved in mitigating the innate immune response at days 1, 3 and 7 post-infusion across the infarct demonstrating less pathological markers at both days 3 and 7 post infusion in the spatial data, corroborating the findings from the corresponding snRNAseq dataset.

We recently showed that a decellularized myocardial ECM hydrogel elicits pro-reparative effects in subacute and chronic MI, and regional bioactivity of the locally administered ECM biomaterial^[11e]^. Derived from the liquid ECM hydrogel, the infusible ECM is comprised of the hydrogel’s liquid precusor without the larger particulates. Here, even when administering the infusible version of the ECM biomaterial, we still measure significant pro-reparative effects, yet at earlier timepoints in MI pathophysiology. Compared to the ECM hydrogel, acute MI treatment with the infusible ECM demonstrated starker mitigation of the innate immune response, anti-inflammatory macrophage polarization, and promotion of MHC-II APCs, while the ECM hydrogel demonstrated preservation of the resident macrophage population and T-cell polarization. Similar to the ECM hydrogel studies, the iECM’s spatial samples also corroborated these acute pro-reparative responses found with snRNAseq, demonstrating that snRNAseq and spatial transcriptomics gain effective power when they are cross-validated. Finally, these subtle pro-reparative differences between the ECM hydrogel and infusible ECM could be due to the higher molecular weight components lost in iECM manufacturing, providing rationale towards understanding what key components of ECM biomaterials enhance pro-repair in acute MI.

The iECM has shown predominant effects in modulating the immune response as early as 1 day post-MI and post-infusion. Monocytes, largely depleted 1 day post-MI, demonstrated regulation of the innate immune response and anti-inflammatory macrophage polarization. In addition, APCs exhibited strong activationof MHC class II genes. While all nucleated cells display varying levels of MHC molecules to interact with antigens, MHC-II^high^ immune cells are implicated in leukocyte activation, which can exacerbate inflammation and vascular permeability following an ischemic insult^[48a]^. MHC-II^high^ immune cells and macrophages are closely associated with the inflammatory phase of the early days of ischemia, with low expression of MHC-II molecules signalling in the reparative phase and attenuation of inflammation^[48c]^. Finally, among macrophages, subsetted macrophages exhibited transcriptomic shifts towards an anti-inflammatory macrophage phenotype at 1 day post-infusion, which was sustained at days 3 and 7 post infusion after iECM degradation, demonstrating the downstream effects of iECM administration in MI.

Given that the iECM degrades at 3 days post-infusion, we also noted that the iECM is also able to induce pro-repair responses at both days 3 and 7 post-infusion. With that in mind, at day 7, saline treated cell types have a mixed expression profile with markers of pro-repair with detrimental markers, such as reduced angiogenesis, heightened inflammation, and increased apoptosis compared to days 1 and 3 post infusion. For instance, day 7 lymphatic endothelial cells had some pro-reparative gene expression via *Efnb2^[31]^,* a marker of the normal reparative reaction post-MI and upregulation of pro-apoptotic *Igfbp5^[32]^*. In addition, day 7 mural cells exhbited *Igf1^[38]^*, a marker of cardioprotection, and *Tmod3^[39]^*, a marker of reduced angiogenesis. Finally, day 7 fibroblasts demonstrated both pro-reparative (*Pde3a*) and pro-fibrotic (*Tgfbr3*) markers, showing that the overall injury and downstream fibrotic response continue post iECM administration. While pro-repair markers are sustained up to 7 days post-infusion, this presents rationale for multiple dosing studies to maintain iECM-induced pro-repair at later timepoints.

## 4. Conclusion

With all this in mind, these results suggest that the iECM can modulate each cell type’s bioactivity in the infarcted heart and allow for an overall pro-reparative response in treating MI. Uniquely, iECM can thus further modulate the immune response of infiltrating monocytes, and thus promoting macrophage polarization because of intravascular delivery at acute timepoints. Similar to the ECM hydrogel, by modulating the immune response, promoting fibroblast activation, increasing vasculature development in the endothelium, lymphatics, and smooth muscle cells, eliciting cardiomyocyte development and salvage, and finally promoting neuronal cell development, the iECM can thus lead to overall mitigation of negative LV remodeling and improve cardiac function.

## 5. Experimental Section

### 5.1. iECM Preparation and Characterization

The infusible ECM (iECM) was prepared from porcine LV myocardial tissue and characterized as mentioned previously^[12a]^. Similar to ECM hydrogel preparation, the tissue was chopped, decellularized, lyophilized, partially enzymatically pepsin digested in HCl, then buffered. The liquid ECM was then subjected to high centrifugation, where the supernatant was extracted and isolated, denoting the iECM. The iECM was then dialyzed, passed through 0.22 μm syringe filters, and lyophilized and stored at -80°C for future use. Before intracoronary administration, the iECM was then resuspended in sterile water to a final concentration of 10 mg/mL for at least 30 minutes at room temperature, as demonstrated previously^[12a]^ .

### 5.2. Surgical Procedures

All procedures in this study were approved by the Committee on Animal Research at the University of California San Diego and in accordance with the guidelines of the Association for the Assessment and Accreditation of Laboratory Animal Care (A3033-01). All surgeries were performed using aseptic conditions. All animals used in the study were adult female Sprague Dawley rats (225 to 250 g). To induce MI, all animals underwent ischemia–reperfusion surgery to occlude the left main artery for 35 min as previously describeds^[9a, 12a]^. Animals were administered either iECM or saline (200 μL) via intracoronary infusion 10 minutes post reperfusion. Intracoronary infusion was induced through clamping of the aorta with 200 μL of iECM or saline injected into the LV lumen via a 30G needle as previously described^[12a]^.

### 5.3. Tissue Processing

All animals were harvested at 1, 3, or 7 days post infusion. Half of the animal hearts per each model group were used were cut into 6 slices using a stainless-steel rat heart slicer matrix (Zivic Instruments) with 1.0 mm coronal spacing. Odd slices were frozen in TissueTek OCT^TM^ and sectioned into 10 μm thick slices and placed onto a 10X Visium Spatial Transcriptomics Slide or a regular histology slide. Tissue sections adjacent to the Visium section were taken within 100 μm for IHC. For the other half of hearts, the LV free wall was isolated from even slices of tissue and flash frozen in liquid nitrogen to preserve RNA as previously described. The remaining slices were fresh frozen in TissueTek OCT freezing medium for histology.

### 5.4. Nuclei Isolation and snRNAseq

Nuclei from cardiac tissue was isolated as mentioned previously. In brief, flash frozen LV free wall samples were resuspended in nuclei lysis buffer (Millipore Sigma, Nuclei EZ prep, NUC101), 0.2 U/μL RNase inhibitor (Enzymatic Y9240L) and finely chopped with scissors. Samples were homogenized with a 2 mL dounce grinder (KIMBLE) with the lysates filtered through 100 μm, 50 μm, 20 μm strainers (CellTrics filters), respectively. The samples were centrifuged at 1000g for 5 min at 4°C to pellet nuclei. The nuclear pellet was resuspended in a sucrose gradient buffer and centrifuged at 1000g at 4°C for 10 minutes. The pellet was then washed with a nuclei storage buffer containing 10 μg ml−1 4′,6-diamidino-2-phenylindole DAPI and centrifuged at 500 g and 4°C for 5 minutes, with the pellet resuspending in 200 mL of 2% BSA in 1x PBS. Nuclei were then counted on a hemocytometer and volume adjusted to 1000 nuclei/μL for loading. snRNAseq was performed with a microfluidic droplet-based technique provided by 10X Genomics kits (Chromium Next GEM Single Cell 3’ HT v3.1). Quality control for cDNA and libraries were performed on Agilent TapeStation, and library concentrations were determined via Qubit HS DNA kit. Paired-end sequencing was run on a NovaSeq6000 or NovaSeq X Plus instrument. Demultiplexing of sequenced samples were mapped to a rat reference genome (Rnor6.0, including introns). Redundant unique molecular identifiers (UMIs) were eliminated via the Cell Ranger 7.1.0 pipeline from 10X Genomics.

### 5.5. Spatial Transcriptomics

As previously mentioned, harvested OCT-embedded cardiac tissue was cryosectioned onto the fiduciary regions on a 10X Visium slide at 10 μm thickness. Sections were then stained with hematoxylin and eosin, with images obtained on a Nikon Eclipse Ti2-E widefield microscope at 10X magnification. Visium was performed as per manufacturer’s kit instructions (Visium Spatial Gene Expression), with tissue permeabilization time of infarcted rat hearts optimized at 42 minutes. Both protocols utilized barcoding and library preparation, which was validated using an Agilent TapeStation prior to sequencing and quantified via Qubit HS DNA Kit. Paired-end sequencing was done on a NovaSeq6000 or a NovaSeqX Plus instrument. Low-level analysis was performed by mapping to a rat reference genome (Rnor6.0, including introns). The SpaceRanger 1.3 pipeline (10X Genomics) was used to remove redundant UMIs.

### 5.6. Quality Control, Normalization, and Integration

All snRNAseq and Visium data analysis was done using the Seurat package (v5) in R. Both sets of data had raw counts with ribosomal and mitochondrial genes cleaned out and counts adjusted for ambient RNA expression using the SoupX package before normalizing by replciate using log normalization. The FindVariableFeatures method was utilized to find the top 2,000 genes with the highest feature variance. Reference-based integration of the single nuclei datasets was performed via RPCA. The integrated data was then subjected to principal component analysis, and further reduced through uniform manifold approximation and projection (UMAP). The data was then clustered and visualized in UMAP space.

For analyses between iECM and saline treatment groups in snRNAseq, the data was subset by cell type and timpeoint and renormalized using SCTransform by replicate. Integration of all snRNAseq replicates was performed in Seurat to enable harmonized clustering and downstream comparative analyses^18^. Canonical reciprocal correlation analysis (RPCA) was used to determine anchoring cell pairs, and integration anchors were detected through the FindIntegrationAnchors function, utilizing reference-based integration of the single nucleus datasets via RPCA. Once the iECM and saline datasets were integrated together, the integrated set was then subjected to principal component analysis, and further reduced through uniform manifold approximation and projection (UMAP) to subject the data into a consensus UMAP space. The data was then clustered and visualized in UMAP space. All canonical cell types (macrophages, endothelial cells, cardiomyocytes, fibroblasts, neuronal cells, smooth muscle cells, and lymphatic endothelial cells) were identified through literature review of the gene signatures^39^ and using the FindAllMarkers function in Seurat v4 (P_adj_ < 0.05, logFC > 0.25, min.diff.pct > 0.25, assay = RNA). Spatial transcriptomic analyses of ECM and saline treatment groups were subjected to the same integration process to ensure comparisons were done in a consensus UMAP space.

For each relevant cell type, the data were subsetted through novel gene markers, integrated via RPCA, renormalized via SCTransfomr, subjected to principal component analysis, and reduced through UMAP to subject each cell type across treatment condition into a consensus UMAP space.

All analyses between iECM and saline were conducted using the FindMarkers function in Seurat v4 (P_adj_ < 0.05, logFC > 0.25, min.pct > 0.25, assay = SCT [snRNAseq] or assay = Spatial [Visium]).

### 5.7. Gene Ontology Enrichment

Gene ontology (GO) was performed using the gene set enrichment analysis software (UC San Diego and Broad Institute). Inputs for GO analyses were determined using the FindMarkers function in the R package Seurat (P_adj_ < 0.05, logFC > 0.25, assay = SCT [snRNAseq] or assay = Spatial [Visium]) to find differential genes between ECM and saline cells.

### 5.8. Immunohistochemistry

After euthanasia, odd sections of the heart were embedded in OCT for cryosectioning. Prior to running the 10X Visium Spatial Transcriptomics protocol, slides were stained with H&E and scanned at 20X using an Olympus VS120 Slide Scanner. All hearts were sectioned at 10 um thickness. For all histology and immunohistochemistry, slides were imaged at 20X using an Olympus VS200 slide scanner (Evident Scientific, Walthan, MA). All histological and immunohistological (IHC) assessment was performed by investigators who were blinded to each group. To visualize infarct size and tissue morphology, heart sections were stained with hematoxylin and eosin (H&E). Infarcts were manually traced to quantify infarct size for animals that were utilized for snRNAseq and spatial transcriptomics.

## Contributions

J.M.M., A.C., Q.L., M.B.N, K.R.K., and K.L.C. designed and performed the experiments, analyzed the data, and wrote the manuscript. J.M.M., A.C., Q.L., J.Y., V.K., M.L.K. Q.L., E.G., E.W., and S.N. performed the snRNAseq and Visium experiments and analyzed the data. A.C. and M.B.N. performed quality control on iECM batches. Z.F. helped with single nucleus RNAseq and Visium experiments. C.G.L. performed rat surgeries and intracoronary infusions. K.L.C. conceived the project and provided funding. All authors reviewed the results and commented on the manuscript.

## Acknowledgements

This research was funded in part by the National Institutes of Health National Heart, Lung, and Blood Institute (NHLBI) (R01HL165232 to KLC). J.M.M. was supported by the National Institutes of Health (NIH), NHLBI Training Grant T32HL105373. A.C. was supported by the NIH, NIBIB Training Grant T32EB009380. J.M.M (23PRE1023221) and A.C. (24PRE1180449) were supported by an American Heart Association pre-doctoral fellowship. M.B.N was supposed by an American Heart Association post-doctoral fellowship (24POST1242447). M.L.K was supported by the NSF GRFP. K.L.C. is a cofounder, board member, consultant for, and holds equity interest in and receives income from Ventrix Bio, Inc. All other authors have reported that they have no relationships relevant to the contents of this paper. This work also utilized the UC San Diego IGM Genomics Center utilizing an Illumina NovaSeq 6000 and NovaSeq X that was purchased with funding from a National Institutes of Health SIG grant (#S10 OD026929), and the UCSD School of Medicine Microscopy Core grant (P30 NS047101).

## Data Availability Statement

Single nucleus RNA sequencing and spatial transcriptomic sequencing datasets have been deposited to the Gene Expression Omnibus and are available under accession GSE299772. All other data supporting the findings in this study are included in the main article and associated files.

## Supporting Information

See corresponding document.

