## Supplementary Information for "Infusible Extracellular Matrix Biomaterial Enhances Cell-Specific Pro-Repair Responses Following Acute Myocardial Infarction"

A. Chen

Program in Materials Science and Engineering, Sanford Consortium for Regenerative Medicine, University of California San Diego, La Jolla, California 92037, United States

J. Yu, V.K. Ninh, Z. Fu, K.R. King

Shu Chien-Gene Lay Department of Bioengineering, School of Medicine, University of California San Diego, La Jolla, California 92037, United States

K. L. Christman

Shu Chien-Gene Lay Department of Bioengineering, Program in Materials Science and Engineering, Sanford Consortium for Regenerative Medicine, Sanford Stem Cell Institute, University of California San Diego, La Jolla, California 92037, United States

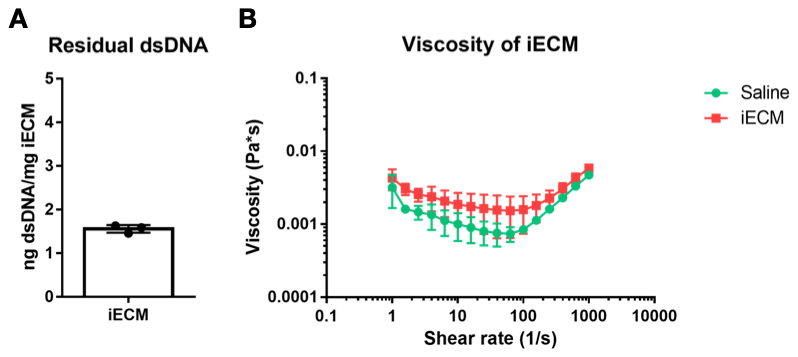

**Figure S1. iECM Quality Control Data**. **A)** Double stranded DNA (dsDNA) content of the iECM. Sample size: n = 3. **B)** The viscosity of the iECM relative to saline. Sample size: n = 4.

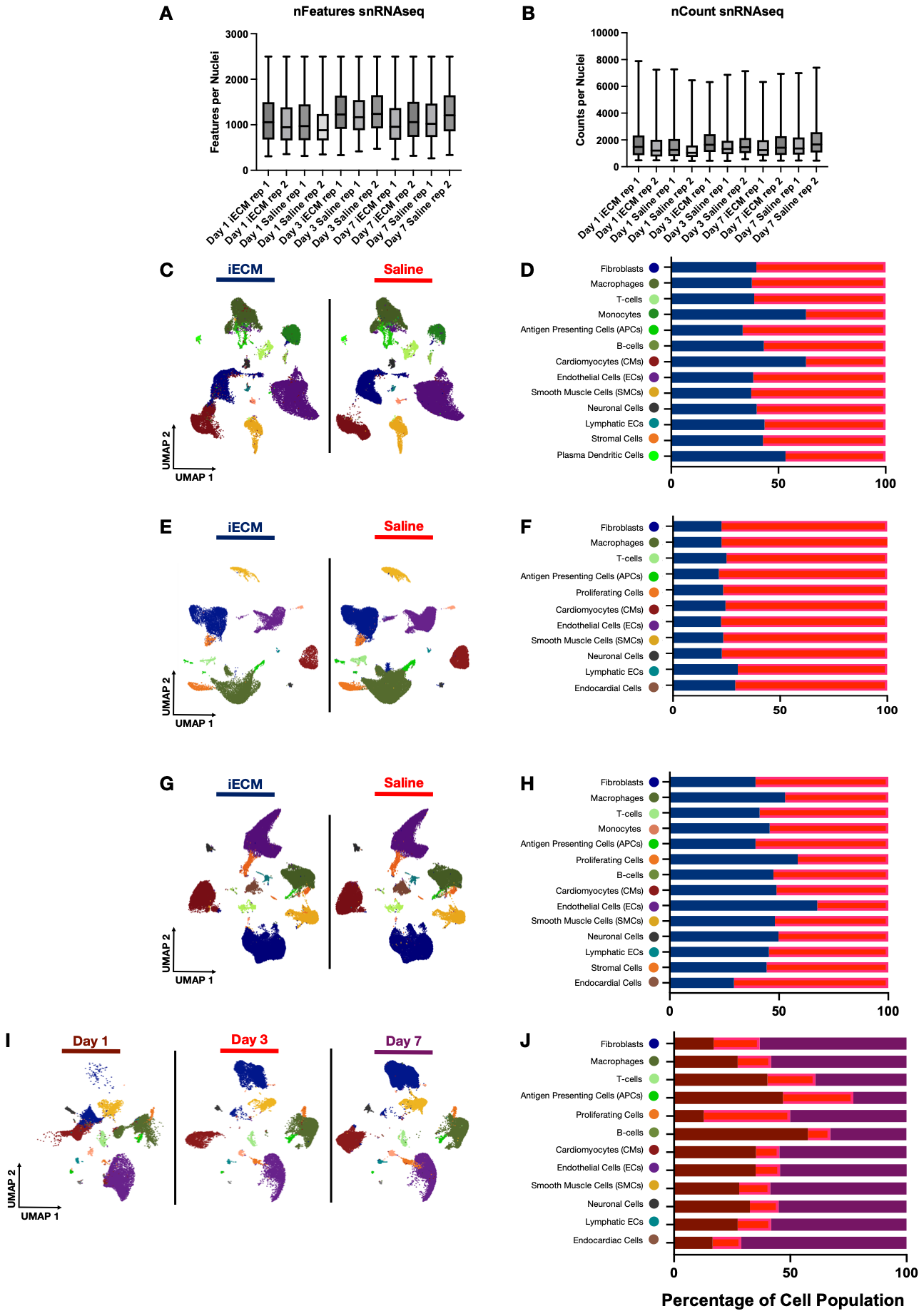

**Figure S2. Single Nucleus RNA Sequencing Quality Control and Cell Type Overview**. **A-B)** Quality metrics of samples and replicates for single nucleus RNA sequencing (snRNAseq) samples represented in features per nuclei (nFeatures) and genes per sample (nCounts). Data are presented as box and whisker plots. **C-D)** Single nucleus RNAseq was performed on infusible ECM and saline hearts harvested 1 day post infusion, where coarse clustering **(C)** defined primary cell types found in the heart with relative percentages and total number of cells per each primary cell time and per treatment **(D). E-F)** Single nucleus RNAseq was performed on infusible ECM and saline hearts harvested 3 day post infusion, where coarse clustering **(E)** defined primary cell types found in the heart with relative percentages and total number of cells per each primary cell time and per treatment **(F). G-H)** Single nucleus RNAseq was performed on infusible ECM and saline hearts harvested 7 day post infusion, where coarse clustering **(G)** defined primary cell types found in the heart with relative percentages and total number of cells per each primary cell time and per treatment **(H). I-J)** Single nucleus RNAseq was performed on infusible ECM harvested at 1, 3 and 7 days post infusion, where coarse clustering **(I)** defined primary cell types found in the heart with relative percentages and total number of cells per each primary cell time and per treatment **(J).** Sample size: n = 4 pooled into n = 2 technical replicates for iECM and saline except Day 3 iECM with n = 2 pooled into n = 1 technical replicate. Day 1: iECM = 33661 cells, saline = 44949 cells; Day 3: iECM = 19526, saline = 64869, Day 7: iECM = 57048 cells, saline = 56229 cells.

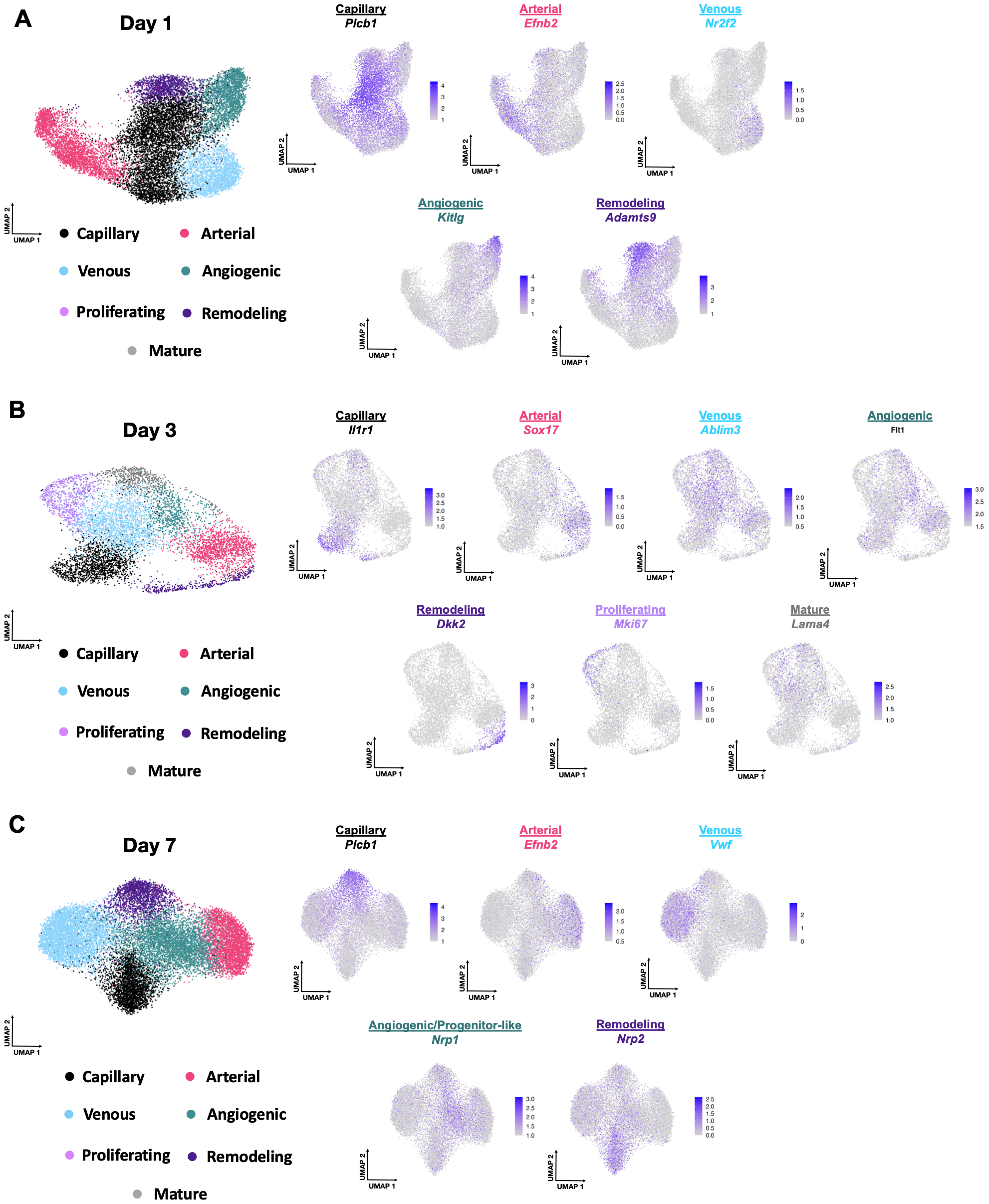

**Figure S3. The Endothelial Cell Subpopulation Phenotype Breakdown Across Time. A-C)** Day 1 **(A)**, Day 3 **(B)**, and Day 7 **(C)** cluster subpopulations identified.

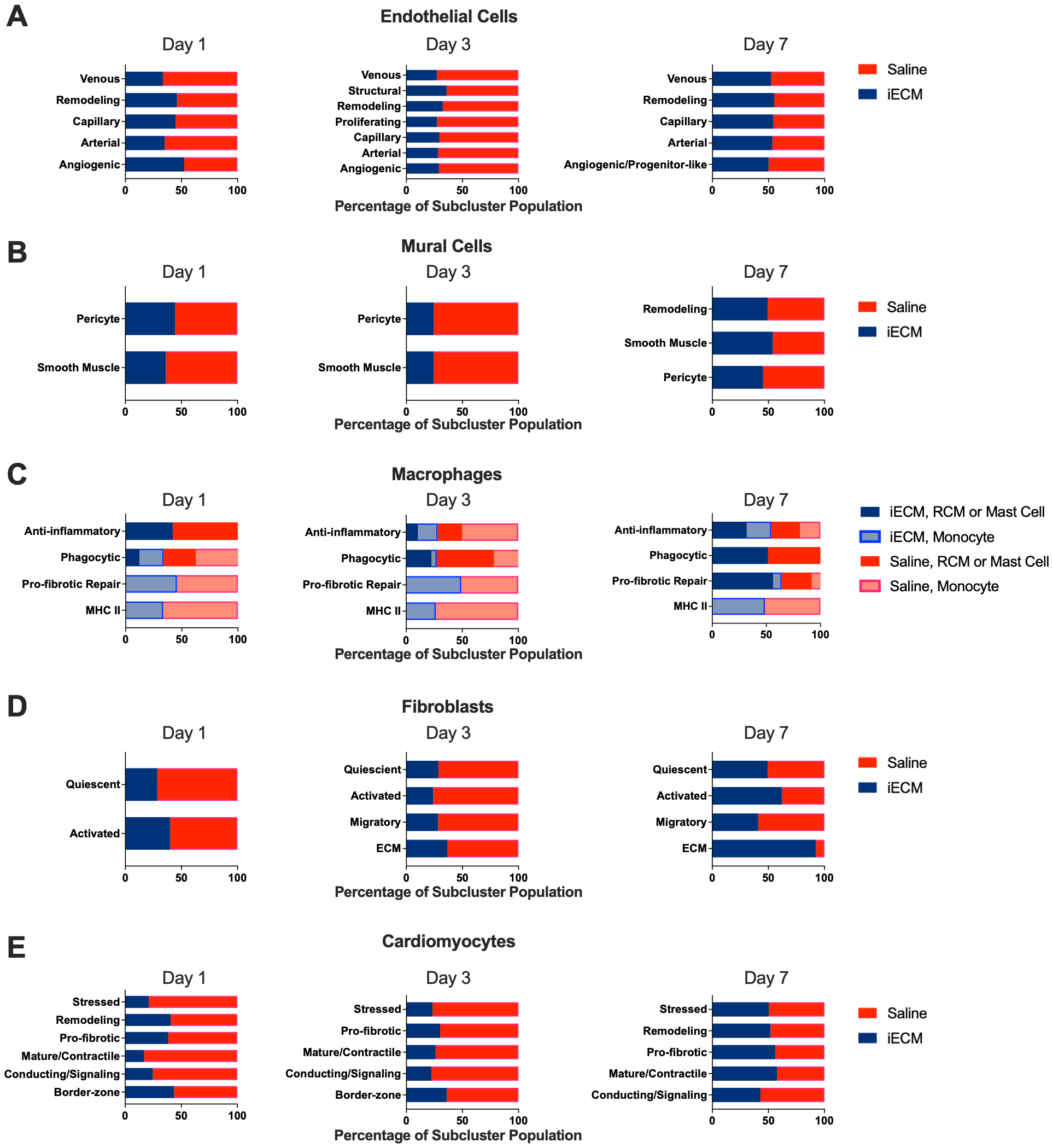

**Figure S4. Relative Subcluster Cell Proportions Between Treatment Groups. A-E)** Endothelial (A), mural (B), macrophage (C), fibroblasts (D), cardiomyocyte (E) subcluster proportions defined by either iECM (blue) or saline (red) treatment at days 1, 3, and 7 post infusion. Macrophages are further divided by iECM, resident cardiac macrophages (RCM) or mast cell origin (dark blue), iECM, monocyte origin (light blue), saline resident cardiac macrophages (RCM) or mast cell origin (red), or saline, monocyte origin (light red).

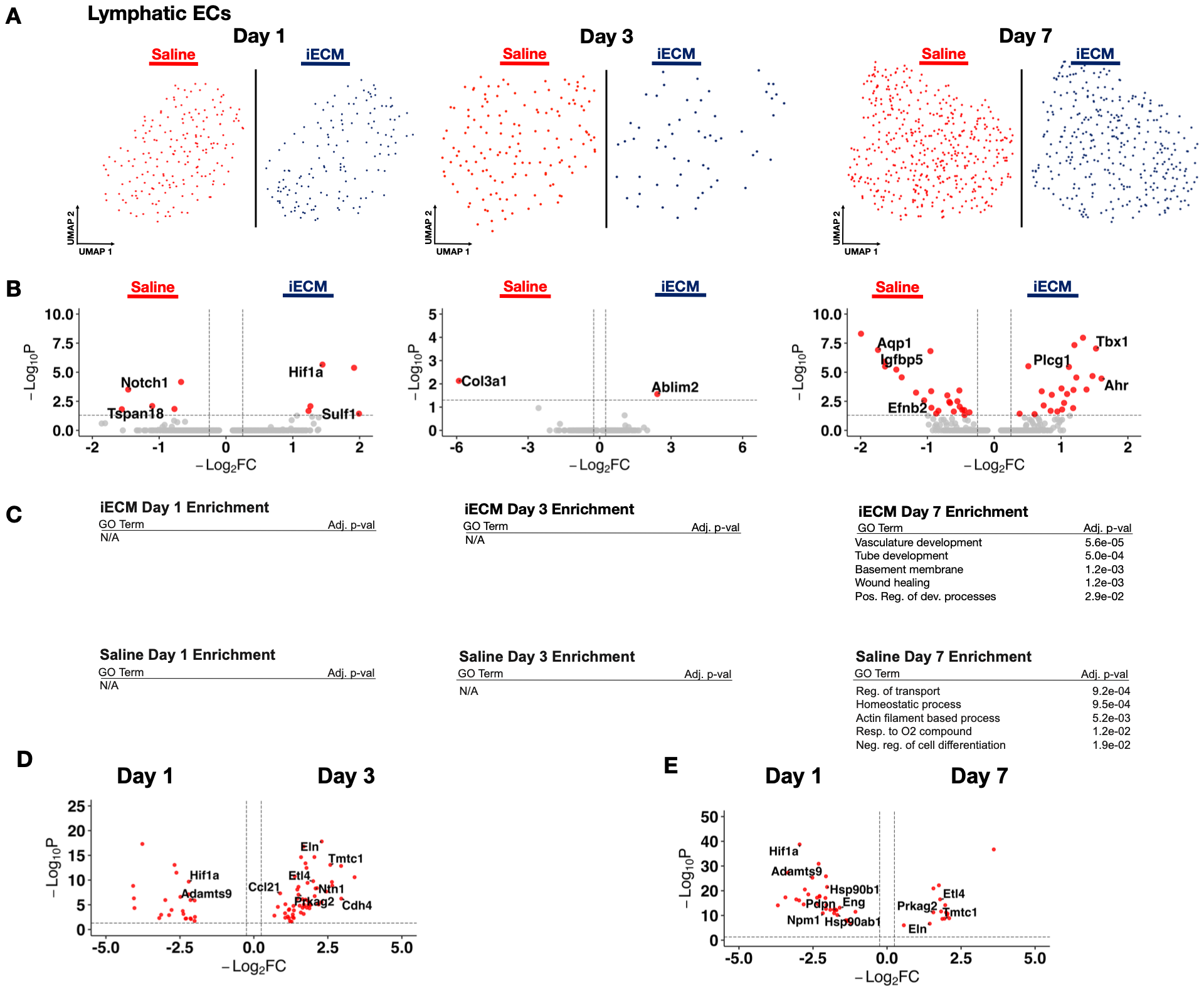

**Figure S5. iECM Promotes Lymphatic Endothelial Cell Development. A)** UMAP of lymphatic endothelial cells in iECM and saline treated samples split across days 1, 3 and 7 post infusion. **B)** Differentially expressed genes in lymphatic endothelial cells from iECM and saline samples at days 1 and 7 post-infusion. **C)** All iECM and saline specific differentially expressed genes were subjected to GO enrichment. **D)** Differentially expressed genes of iECM treated samples at days 1 and 3 post infusion. **E)** Differentially expressed genes of iECM treated samples at days 1 and 7 post infusion.

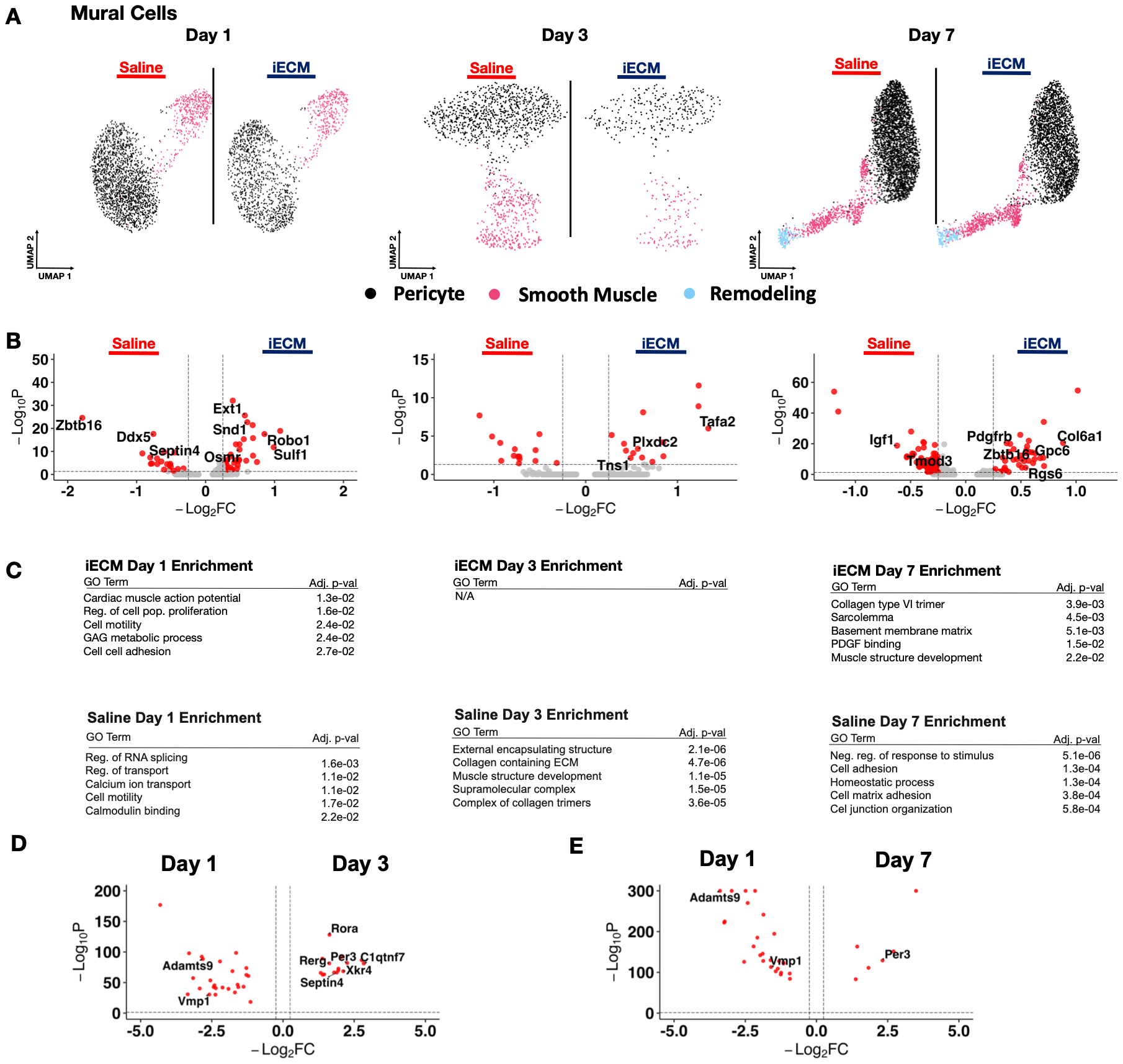

**Figure S6. iECM Promotes Mural Cell Development. A)** UMAP of mural cells in iECM and saline treated samples across days 1, 3 and 7 post infusion. **B)** Differentially expressed genes of iECM and saline mural cells at days 1 and 7 post-infusion. **C)** All iECM and saline specific differentially expressed genes were subjected to GO enrichment. **D)** Differentially expressed genes of iECM samples at days 1 and 3 post infusion. **E)** Differentially expressed genes of iECM samples at days 1 and 7 post infusion.

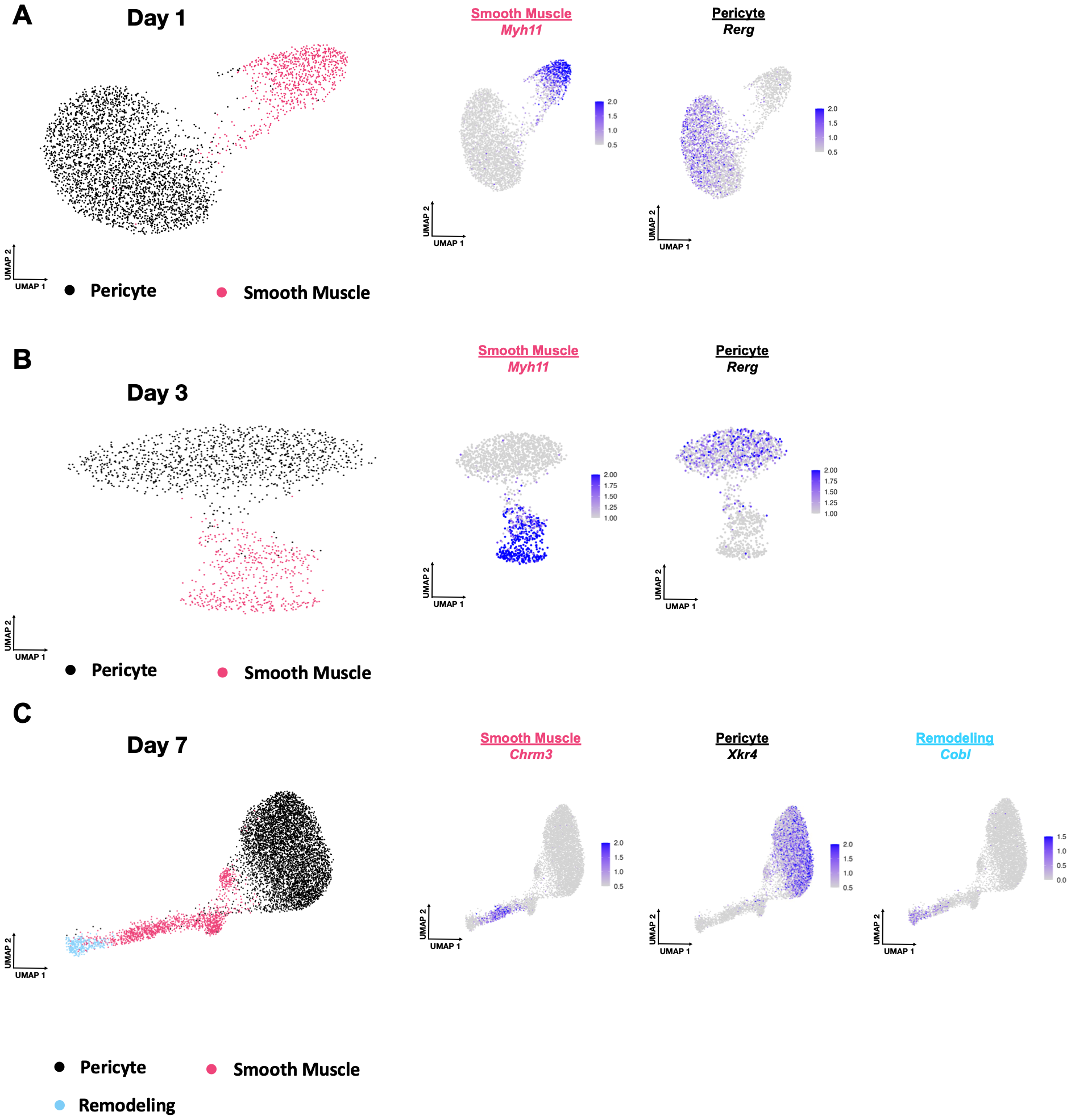

**Figure S7. Marker Genes for Novel Mural Cell Populations. A)** Marker genes for the pericyte and smooth muscle cell subclusters found by subclustering mural cells from day 1 post infusion. **B)** Marker genes for the pericyte and smooth muscle cell subclusters found by subclustering mural cells from day 3 post infusion. **C)** Marker genes for the pericyte, smooth muscle cell, and remodeling subclusters found by subclustering mural cells from day 7 post infusion.

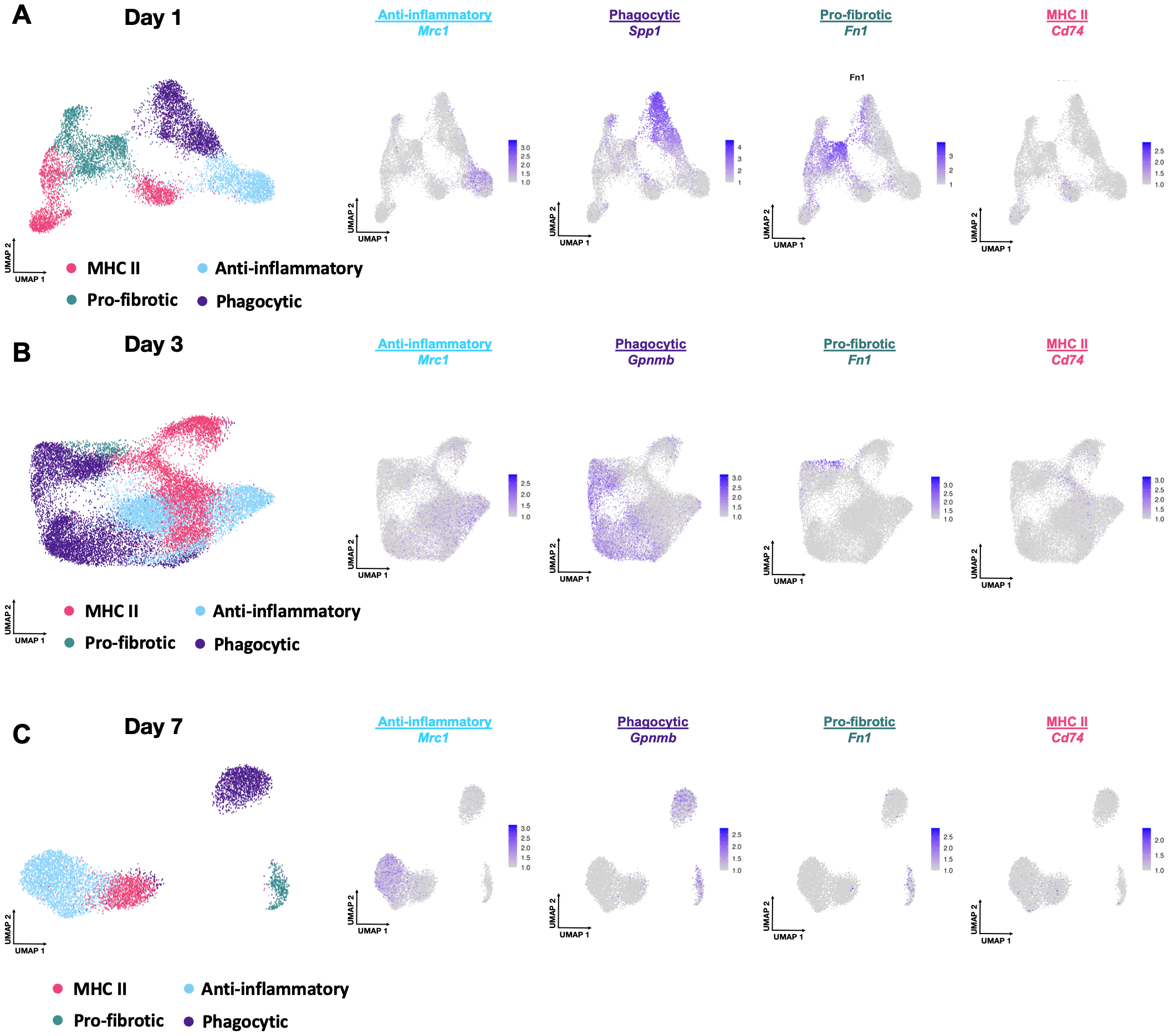

**Figure S8. Marker Genes for Novel Macrophage Populations. A)** Marker genes for the anti-inflammatory, phagocytic, pro-fibrotic and MHC II subclusters found by subclustering macrophages from day 1 post infusion. **B)** Marker genes for the anti-inflammatory, phagocytic, pro-fibrotic and MHC II subclusters found by subclustering macrophages from day 3 post infusion **C)** Marker genes for the anti-inflammatory, phagocytic, pro-fibrotic and MHC II subclusters found by subclustering macrophages from day 7 post infusion.

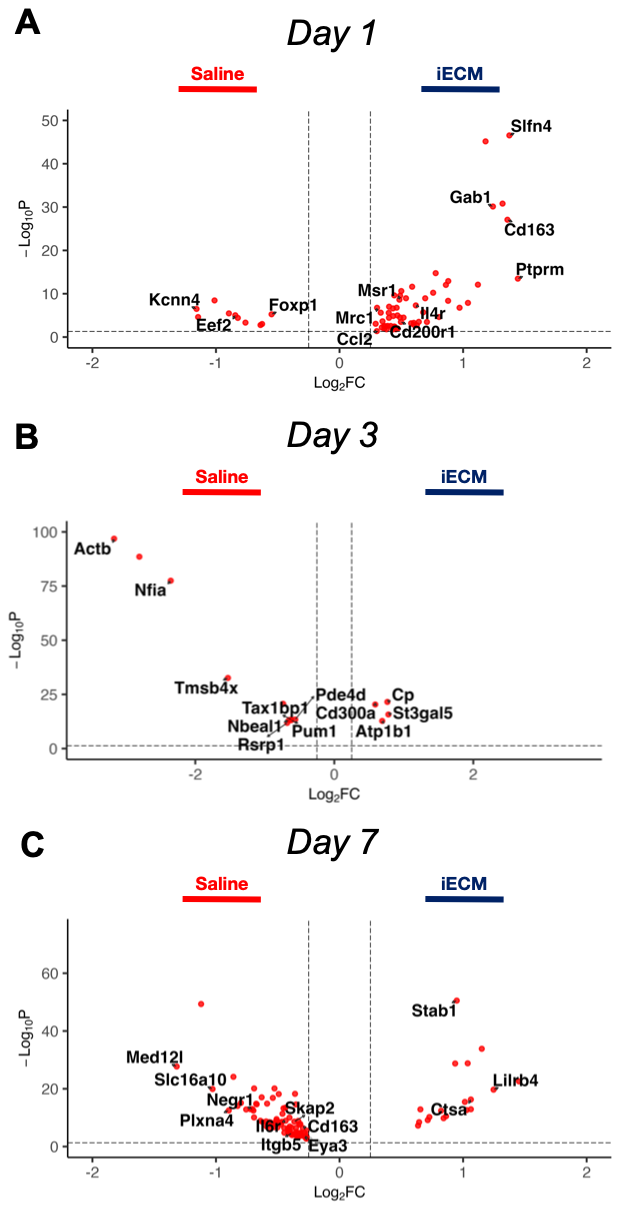

**Figure S9. Differentially Expressed Genes in Anti-Inflammatory Macrophages between iECM and Saline Treatment. A-C)** Volcano plots at days 1, 3, and 7 post infusion of anti-inflammatory macrophages treated with iECM vs. saline.

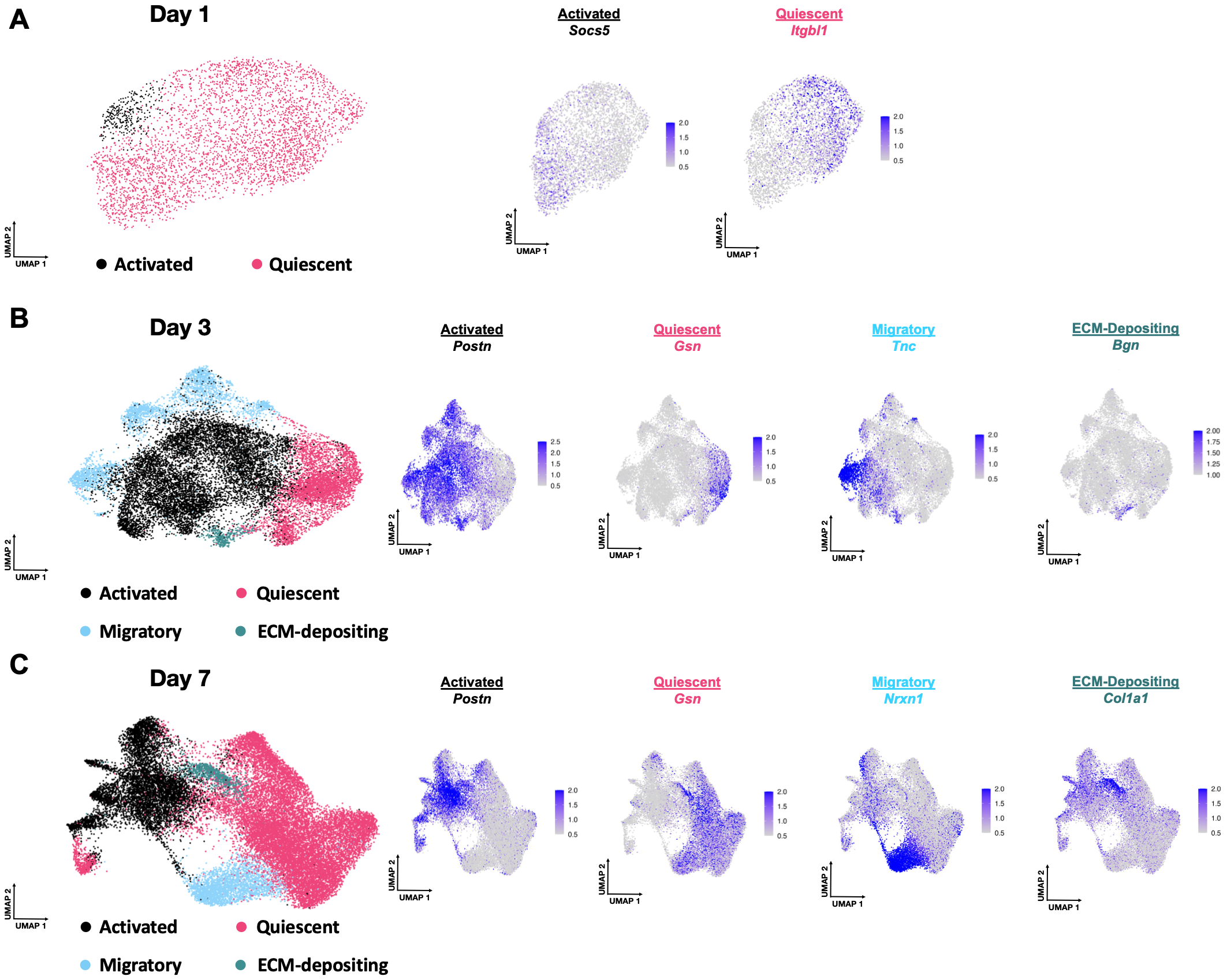

**Figure S10. Marker Genes for Novel Fibroblast Populations. A)** Marker genes for the quiescent, activated, migratory and ECM-depositing subclusters found by subclustering fibroblasts from day 1 post infusion. **B)** Marker genes for the quiescent, activated, migratory and ECM-depositing subclusters found by subclustering fibroblasts from day 3 post infusion. **C)** Marker genes for the quiescent, activated, migratory and ECM-depositing subclusters found by subclustering fibroblasts from day 7 post infusion.

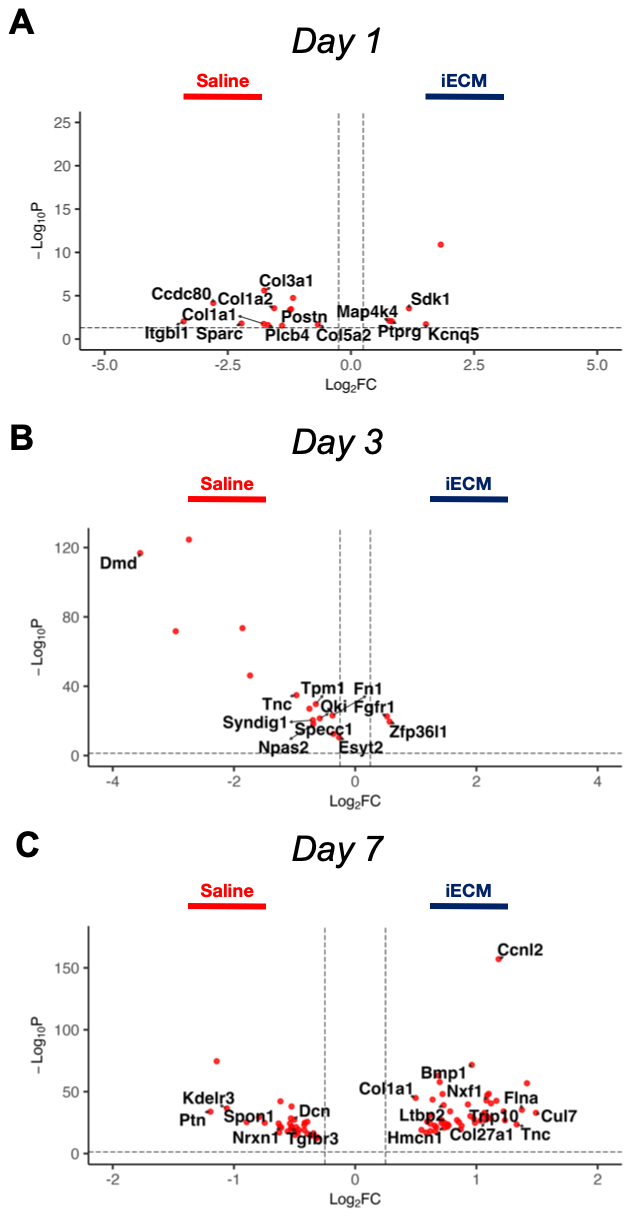

**Figure S11. Differentially Expressed Genes in Fibroblasts between iECM and Saline Treatment. A-C)** Volcano plots at days 1, 3, and 7 post infusion of activated fibroblasts treated with iECM vs. saline.

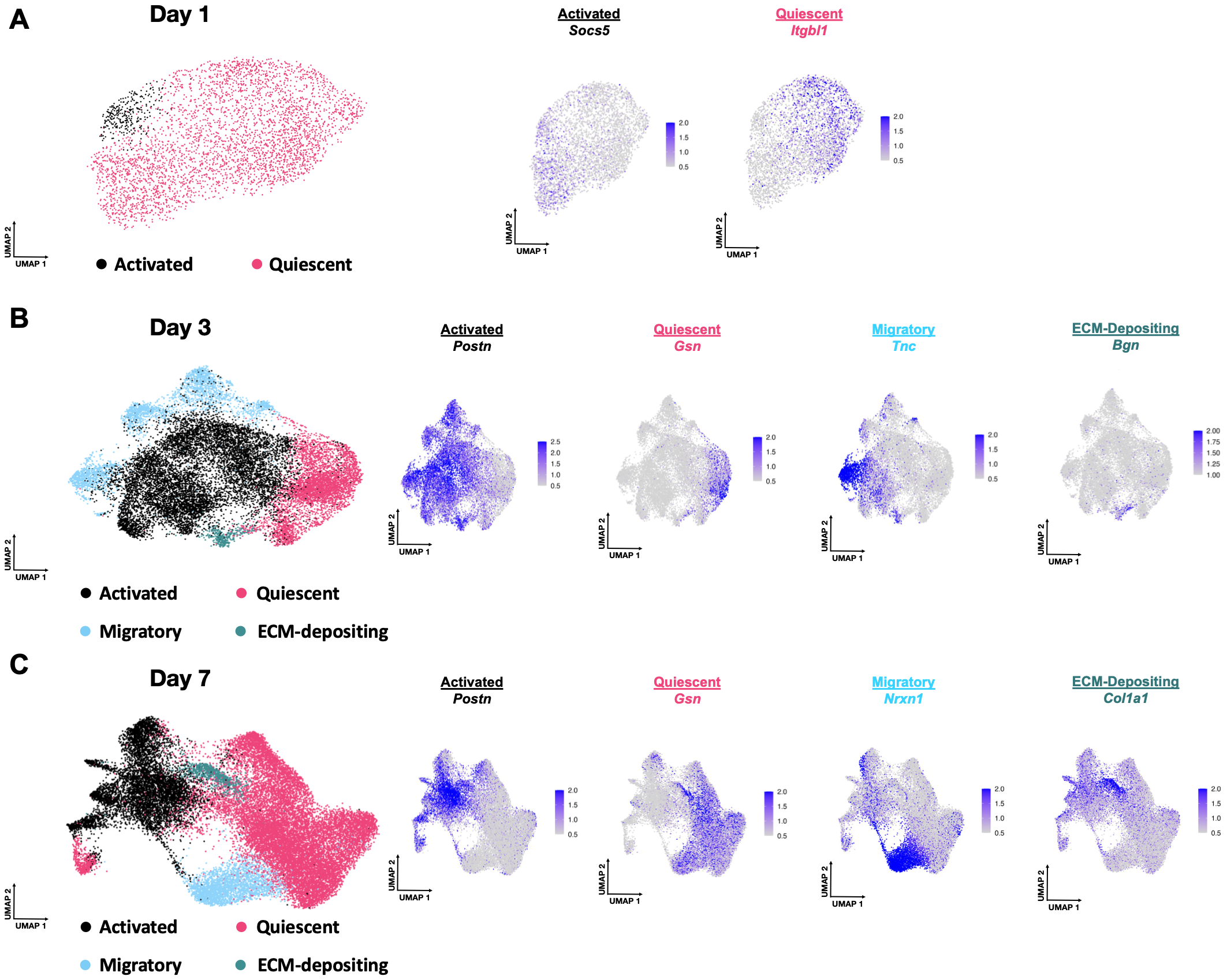

**Figure S12. Marker Genes for Novel Cardiomyocyte Populations. A)** Marker genes for the mature/contractile, conducting/signaling, stressed, pro-fibrotic, border zone and remodeling subclusters found by subclustering cardiomyocytes from day 1 post infusion. **B)** Marker genes for the mature/contractile, conducting/signaling, stressed, pro-fibrotic, and remodeling subclusters found by subclustering cardiomyocytes from day 3 post infusion. **C)** Marker genes for the mature/contractile, conducting/signaling, stressed, pro-fibrotic, and remodeling subclusters found by subclustering cardiomyocytes from day 7 post infusion.

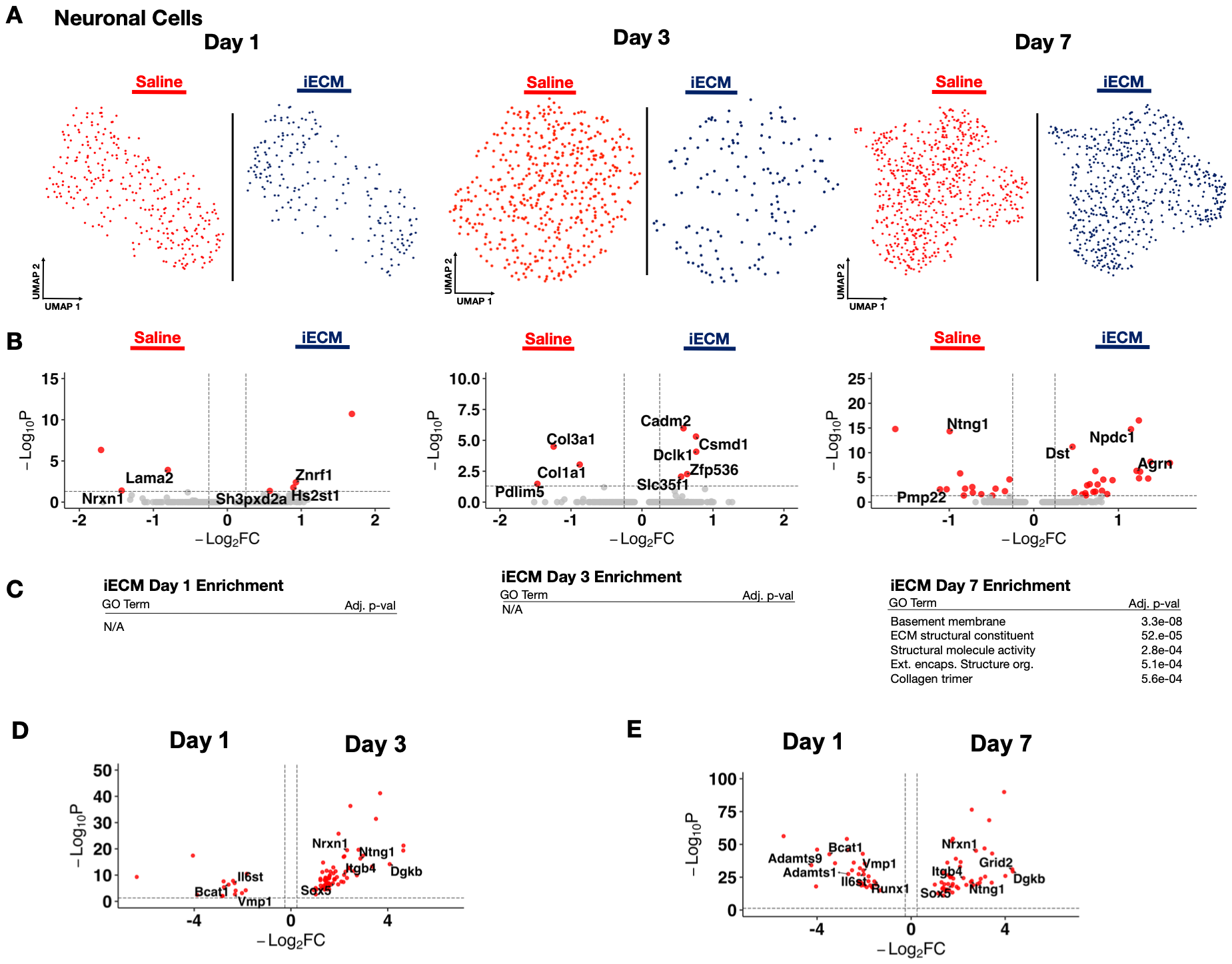

**Figure S13. iECM Promotes Neurogenesis. A)** UMAP of neuronal cells in iECM and saline treated samples split across days 1, 3 and 7 post infusion. **B)** Differentially expressed genes of iECM and saline neuronal cells at days 1, 3, and 7 post-infusion. **C)** All iECM and saline specific differentially expressed genes were subjected to GO enrichment. **D)** Differentially expressed genes of iECM treated neuronal cells at days 1 and 3 post infusion. **E)** Differentially expressed genes of iECM treated neuronal cells at days 1 and 7 post infusion.

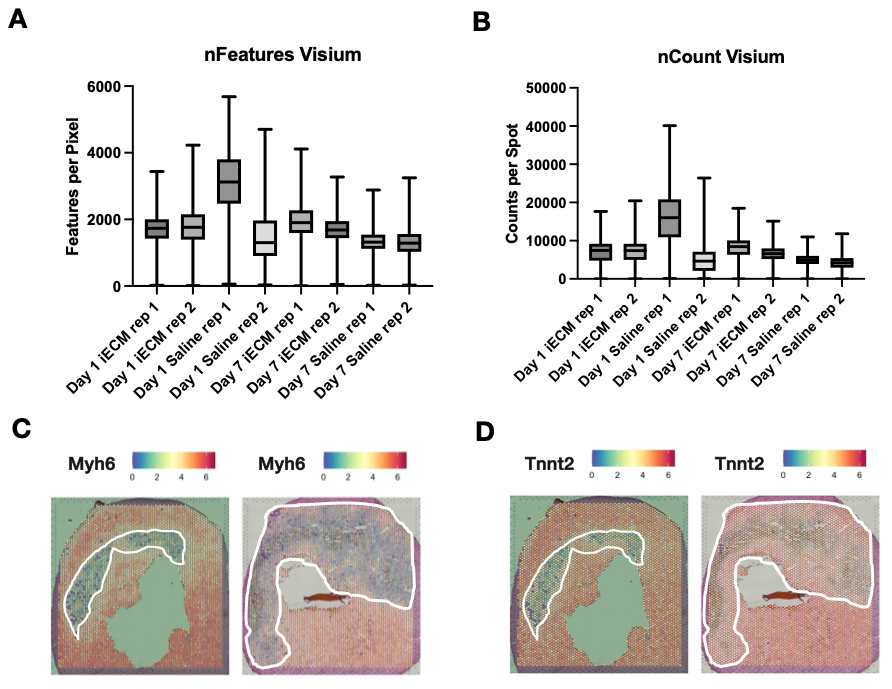

**Figure S14. Visium Quality Metrics and Infarct Zone Strategy. A-B)** Quality metrics of samples and replicates for Visium samples represented in features per pixel (nFeatures) and genes per sample (nCounts). Data are presented as box and whisker plots. **C-D)** A section was used for 10X Visium, with *Myh6* **(C)** and *Tnnt2* **(D)**, two markers for healthy myocardium, measured to segment and identify clusters that are *Myh6* and *Tnnt2* low. White outlines were overlayed onto the coarse clustering plot, indicating which clusters are infarct specific.

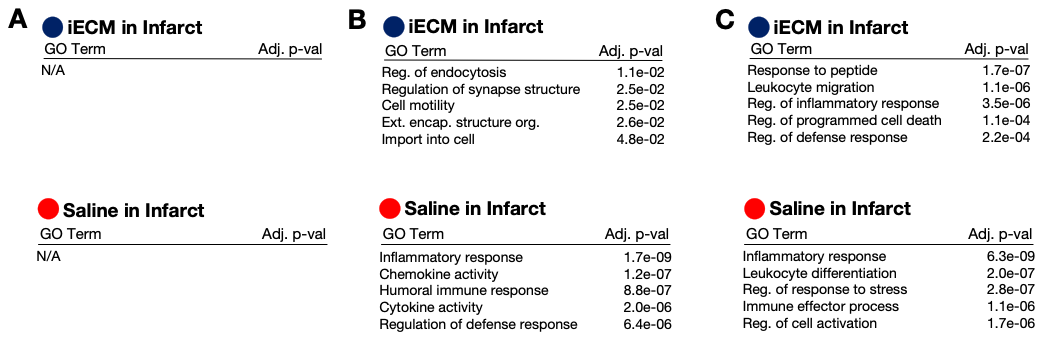

**Figure S15. Gene Ontology for Spatial iECM Samples. A)** Day 1 iECM in infarct (blue) and saline in infarct (red) differentially expressed genes in the iECM and saline treated spatial sections were subjected to GO enrichment. **B)** Day 3 iECM in infarct (blue) and saline in infarct (red) differentially expressed genes in the iECM and saline treated spatial sections were subjected to GO enrichment. **C)** Day 7 iECM in infarct (blue) and saline in infarct (red) differentially expressed genes in the iECM and saline treated spatial sections were subjected to GO enrichment.

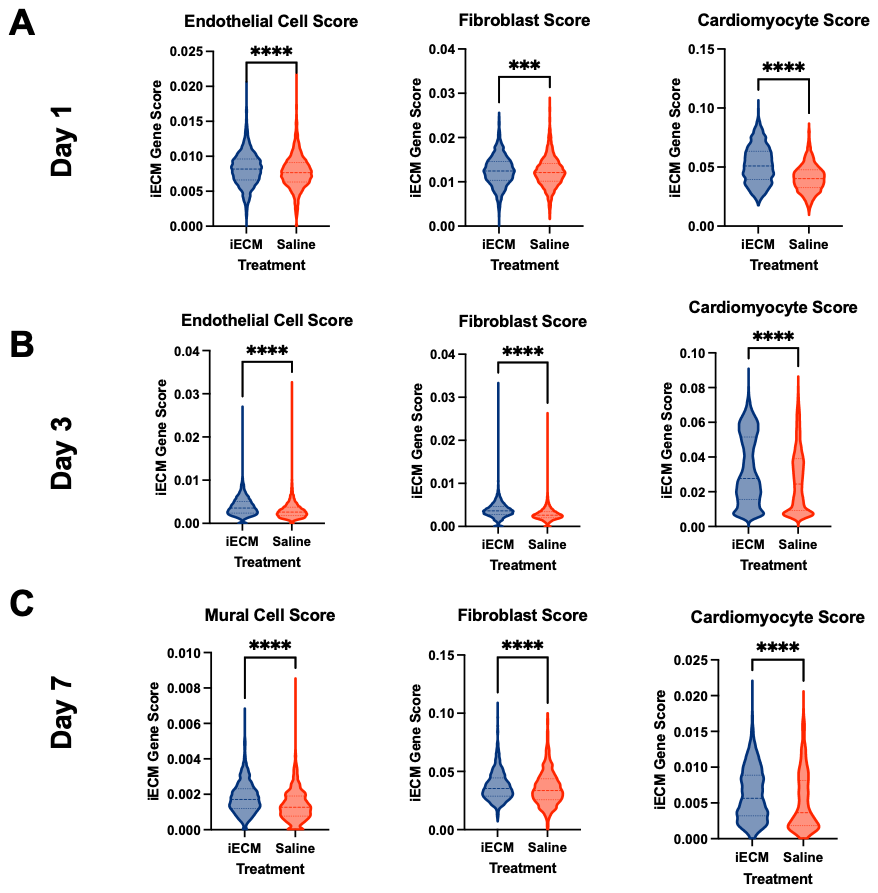

**Figure S16. Infusible ECM Cell-Type Specific Differentially Expressed Genes Mapped onto Spatial Samples. A)** Differentially expressed genes unique to iECM treated endothelial cells, fibroblasts, and cardiomyocytes were scored onto the infarct zone of iECM and saline treated spatial samples from 1 day post-infusion. **B)** Differentially expressed genes unique to iECM treated endothelial cells, fibroblasts, and cardiomyocytes were scored onto the infarct zone of iECM and saline treated spatial samples from 3 day post-infusion. **C)** Differentially expressed genes unique to iECM treated mural cells, fibroblasts, and cardiomyocytes were scored onto the infarct zone of iECM and saline treated spatial samples from 7 days post-infusion. Data are presented as violin plots. Significance was calculated using a Mann-Whitney U-Test. **** P < 0.0001.
